# High accuracy fluorescence-guided cryo-focused ion beam milling

**DOI:** 10.64898/2026.05.11.724418

**Authors:** Davis Perez, Sophia Betzler, Patrick Cleeve, Carmela Villegas, Cali Antolini, Sven Klumpe, Jonathan Schwartz, John M. Mitchels, Shu-Hsien Sheu, Peter D. Dahlberg, Bridget Carragher, David A. Agard, Julia Peukes, Garrett Greenan

**Affiliations:** Biohub, Redwood City, 94065 CA, USA; Ramaciotti Centre for Cryo-Electron Microscopy, Monash University, 15 Innovation Walk, Melbourne, 3800, VIC, Australia; fibsemOS, Melbourne, VIC, Australia; Diamond Light Source, Harwell Science and Innovation Campus, Didcot, OX1 0DE, United Kingdom; Research Institute of Molecular Pathology (IMP), Vienna BioCenter (VBC), Vienna, Austria; Institute of Molecular Biotechnology Austria (IMBA), Vienna BioCenter (VBC), Vienna, Austria; ThermoFisher Scientific, Brno, 62700 Czech Republic; Department of Photon Science, SLAC National Accelerator Laboratory, 2575 Sand Hill Road, Menlo Park, CA 94025, USA; Department of Structural Biology, Stanford University School of Medicine, Fairchild Science Building, 299 Campus Drive West, Stanford, CA 94305, USA; Department of Biochemistry and Biophysics, University of California, San Francisco, San Francisco, CA, USA; Quantitative Biosciences Institute (QBI), University of California, San Francisco, San Francisco, CA, USA

## Abstract

Cryo–electron tomography (cryo-ET) is a powerful approach for visualizing macromolecular structures directly within cells, but its broader application is limited by the difficulty of reliably targeting specific structures during sample preparation. In particular, capturing small or rare objects within cryo–focused ion beam (cryo-FIB) milled lamellae remains a major bottleneck. Here, we present two fluorescence-guided cryo-FIB milling workflows that overcome key sources of targeting error and enable routine capture of structures across a broad range of spatial scales. For larger targets (>500 nm), we present an improved registration-based strategy combining FIB-milled fiducials with depth correction for refractive index mismatch. For smaller targets (150–500 nm), we implement real-time fluorescence guided milling on a commercially available cryo–FIB-SEM platform enabling consistent recovery of small, low-copy organelles such as centrosomes and primary cilia. Both workflows are implemented in a new open-source package fibsemOS and together expand the range of cellular structures accessible to cryo-ET and establish a broadly deployable framework for targeted in situ structural biology.

## Introduction

Cryo-electron tomography (cryo-ET) enables in situ visualization of molecular architecture within cells at nanometer resolution, with subtomogram averaging extending this to near-atomic detail (Nogales and Mahamid, 2024). In cryo-ET, tilt series acquired in a cryo-transmission electron microscope (cryo-TEM) are computationally reconstructed into three-dimensional tomograms, providing a direct view of cellular ultrastructure in its native context. This capability makes cryo-ET uniquely suited for structural cell biology.

A central constraint of cryo-ET is sample thickness: biological specimens must be thinned to 200-300 nm or less to be electron transparent, making it challenging to access and accurately target structures buried deep within cells. For cellular samples several micrometers thick, this necessitates cryo-focused ion beam (cryo-FIB) milling, in which material is removed above and below a region of interest to produce a thin section or ‘lamella’ (Marko et al., 2007; Wagner et al., 2020). In standard workflows, milling is guided by scanning electron microscopy (SEM) and low-current FIB imaging, using surface topology to select regions for thinning (Zachs et al., 2020). While effective for abundant structures, this approach restricts in situ cryo-ET to highly prevalent cellular components such as ribosomes or mitochondria (Kelley et al., 2026) (Fig. 1a).

**Figure 1:**
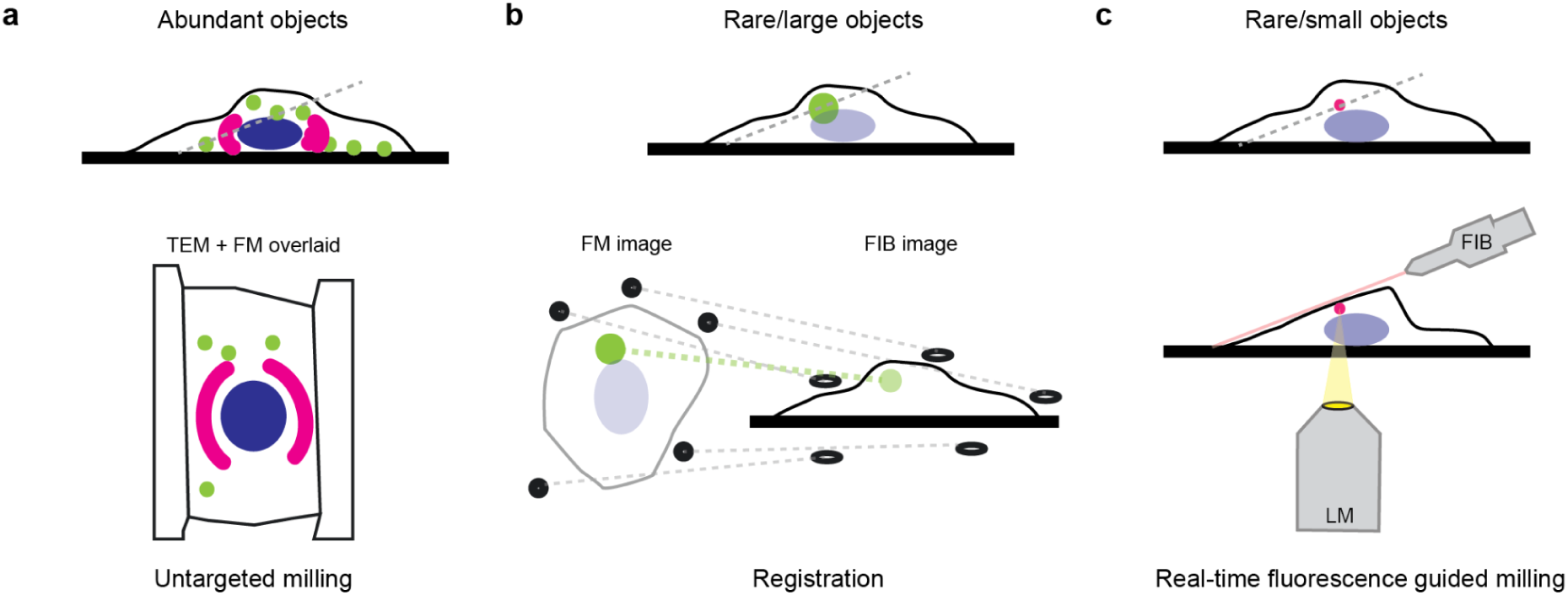
Workflow strategies for LM guided targeting depend on object size and abundance. a) Random milling allows capturing abundant objects by chance. b) For large and rare objects (> 1 μm) registration based approaches allow guiding lamella preparation. Here we introduce an improved registration workflow that allows robustly capturing objects up to 500 nm. c) For rare and small objects (< 500 nm) real-time fluorescence guided milling is needed to capture those reliably. Here we present a robust workflow for capturing objects between 150-500 nm on a commercially available instrument with minor hardware modification.

Precise targeting of small and low-copy-number intracellular structures remains a major bottleneck. One approach is to guide cryo-FIB milling using cryo-fluorescence microscopy, allowing labeled targets to be identified within intact cells. Previous work has demonstrated the promise of this approach (Arnold et al., 2016; Bieber et al., 2021; Fukuda et al., 2014; Gorelick et al., 2019; Klumpe et al., 2021; W. Li et al., 2023; Yang et al., 2026). Most implementations rely on registration-based workflows that map fluorescence coordinates onto the FIB imaging frame (Fig. 1b). These approaches involve correlating fiducials between three-dimensional fluorescence image stacks and two-dimensional FIB images to compute a geometric transformation used for targeting (Arnold et al., 2016). In practice, this process is error-prone: uncertainties in fiducial localization, limitations of the transformation model, sample deformation during milling, and focal shifts arising from refractive index mismatch between cellular material and vacuum (Hell et al., 1993; Petrov and Moerner, 2020; Loginov et al., 2024), collectively produce targeting errors on the order of the lamella thickness or greater. As a result, conventional registration-based targeting is generally limited to micron-scale objects or requires iterative re-registration to achieve acceptable accuracy.

Real-time fluorescence guided milling has recently emerged as an alternative strategy that bypasses many of these limitations (Boltje et al., 2022) (Fig. 1c). In this approach, fluorescence is monitored during milling, so milling can be stopped when signal loss indicates the beam has begun to ablate the object. This semi-destructive strategy has enabled targeting of structures on the order of ∼300 nm (S. Li et al., 2023; Wang et al., 2024), while interferometric approaches have further pushed targeting toward the ∼100 nm scale (Sica et al., 2026). Despite these advances, real-time fluorescence workflows require specialized instrument geometries that allow simultaneous optical and ion beam access to the sample, currently limiting their adoption to very few custom built instruments.

Here, we address the targeting bottleneck through two complementary strategies. First, we establish an accurate registration-based workflow by integrating automated FIB-milled fiducials with a physics-based correction for focal shifts induced by refractive index mismatch. Second, we implement real-time fluorescence-guided cryo-PFIB milling on a commercially available instrument, the Thermo Fisher Scientific Arctis, with minimal hardware modification. To maximize accessibility, both approaches are implemented within fibsemOS, an open-source instrument control framework.

We validate these approaches using fluorescently labeled centrosomes to target individual centrioles, the dominant structural component of centrosomes, as a stringent test case for small, sparse targets. We then extend the real-time fluorescence workflow to target primary cilia, which lack an in situ structural characterization by cryo-ET due to targeting challenges during FIB milling for sample preparation. Together, these results establish robust and potentially broadly accessible strategies for targeting submicron cellular structures, overcoming a key limitation in cryo-ET and expanding its applicability to rare and previously inaccessible biological features.

## Results

### Registration-based targeting

Registration-based correlation is the dominant strategy for fluorescence-guided cryo-FIB targeting, as it is possible with a variety of commercially available cryo-fluorescence microscopes. However, several key sources of error limit its accuracy, and therefore the minimum object size which can be reliably targeted. To address these errors, we developed a workflow which integrates two key advances: First, we incorporated a physics-based depth correction that compensates for focal shifts arising from refractive index mismatch with a tilted interface. Second we incorporated automated placement of FIB-milled fiducials (spot burns) (Martynowycz et al., 2023) across the region of interest, ensuring sufficient control points for an accurate geometric transformation between fluorescence and FIB coordinate systems. Finally, we quantified the targeting accuracy of this workflow and evaluated the remaining sources of error.

Depth correction is central to accurate registration-based targeting. We developed a simulation framework based on vectorial diffraction theory to compute the point spread function (PSF) for an object at a given depth below a tilted interface of refractive index mismatch (Fig. 2a). This calculation is an extension of the work presented in (Hell et al., 1993; Loginov et al., 2024) to the case of a tilted interface (Supplementary Note 1). Since most cryo light microscopes have an optical axis normal to the sample, this interface tilt can either be approximated as 0 for unmilled samples, or taken as the milling angle when the top of the sample above the object has been removed. When unmilled cells are viewed from above, the curved surface acts as a lens, affecting the PSF in unknown ways. For this reason, we choose to prepare a flat, tilted surface above the object and account for this known interface in the simulation.

**Figure 2.**
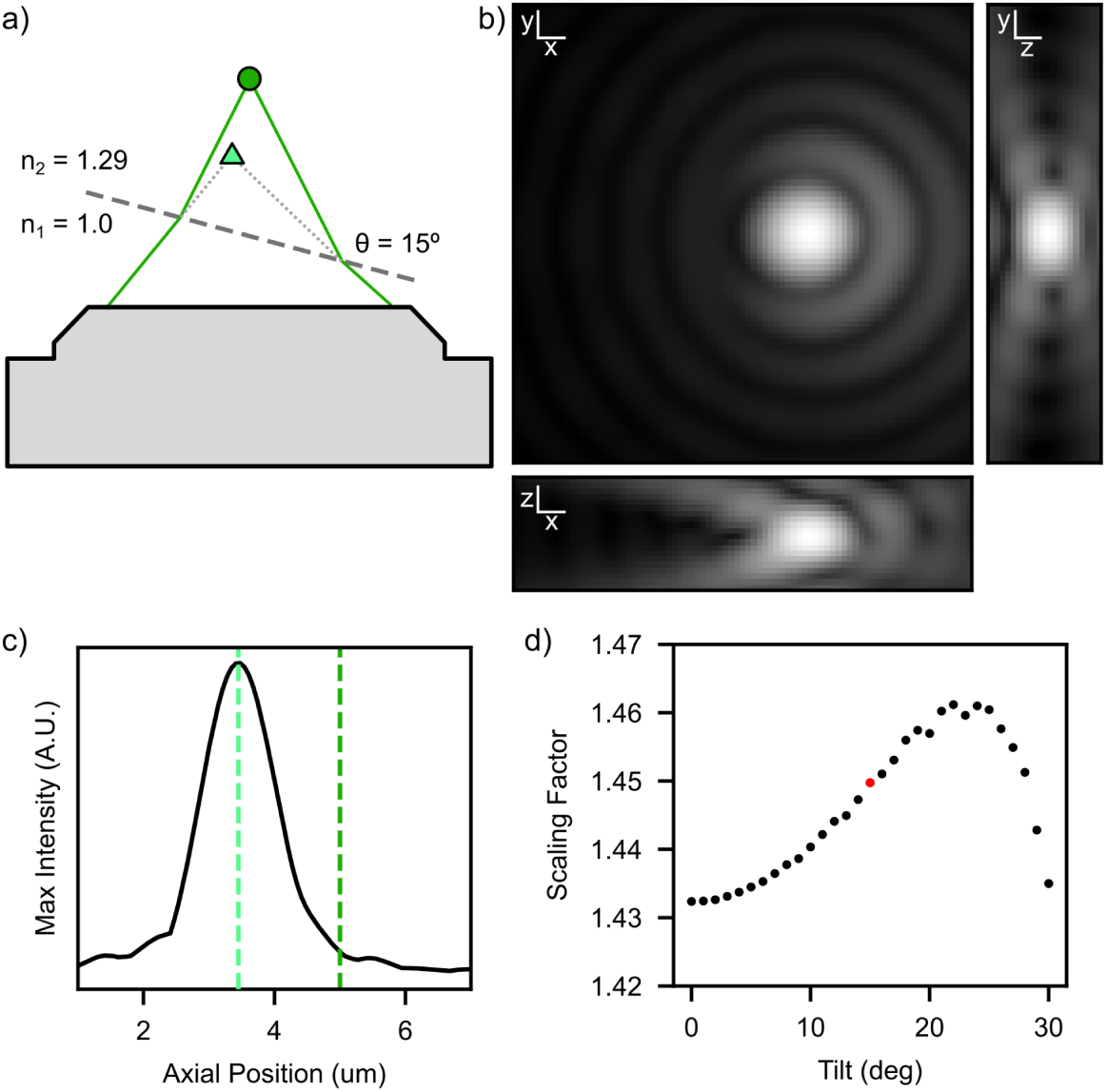
Optics simulation for focal shift from refractive index mismatch. a) Cartoon demonstrating the focal shift effect for conditions relevant to this experiment. The light green triangle and dark green circle show the observed and actual object positions respectively. b) Slices of the PSF calculated for an object 5 μm below an interface at 15° in material with refractive index 1.29 emitting light at 580 nm collected by an objective with NA 0.8. Displayed with gamma 0.3 to show weaker details. c) Axial profile of the PSF shown in (b) constructed by taking the max pixel from each z slice. Dark and light green dashed lines show actual and observed object depth. d) Scaling factors calculated from 31 simulations with the above conditions, varying the interface tilt. The red dot marks the conditions shown in (b) and (c). Scale bars 200 nm x and y, 1 μm z.

Briefly, for a given set of parameters, the calculated PSF is sliced at discrete z-planes to create a simulated z-stack (Fig. 2b). The axial intensity profile is constructed by taking the maximum pixel value from each slice (Fig. 2c). The apparent focal position was defined as the maximum of this profile. Across 352,408 parameter combinations spanning tilt, depth, numerical aperture, wavelength, and refractive index (Table 1), we computed the resulting scaling factors. Representative trends are shown in Fig. 2d and Fig. S2.

**Table 1.** Parameters for optical simulations.

| Parameter | Range start | Range end | Step |
| --- | --- | --- | --- |
| Tilt (degree) | 0 | 30 | 1 |
| Depth (um) | 0.5 | 14.5 | 0.5 |
| NA | 0.2 | 0.9 | 0.1 |
| Wavelength (nm) | 420 | 720 | 50 |
| $n_2$ | 1.22 | 1.46 | 0.04 |

For moderate tilt and depth, the scaling factor varies smoothly with system parameters. At higher tilt or depth, increased aberrations introduce variability due to discretization and peak selection in the axial profile. Despite these effects, the overall dependence remains well behaved within the experimentally relevant regime (<15 μm).

We next experimentally validated the model using a test sample of fluorescent beads embedded in vitrified water droplets (Fig. 3). By measuring the apparent axial shift following removal of a known thickness of material, we directly estimated the scaling factor (Supplementary Note 2). Across multiple samples, we obtained a value of 1.43 ± 0.06, in close agreement with the simulated value of 1.45 for the same conditions.

**Figure 3.**
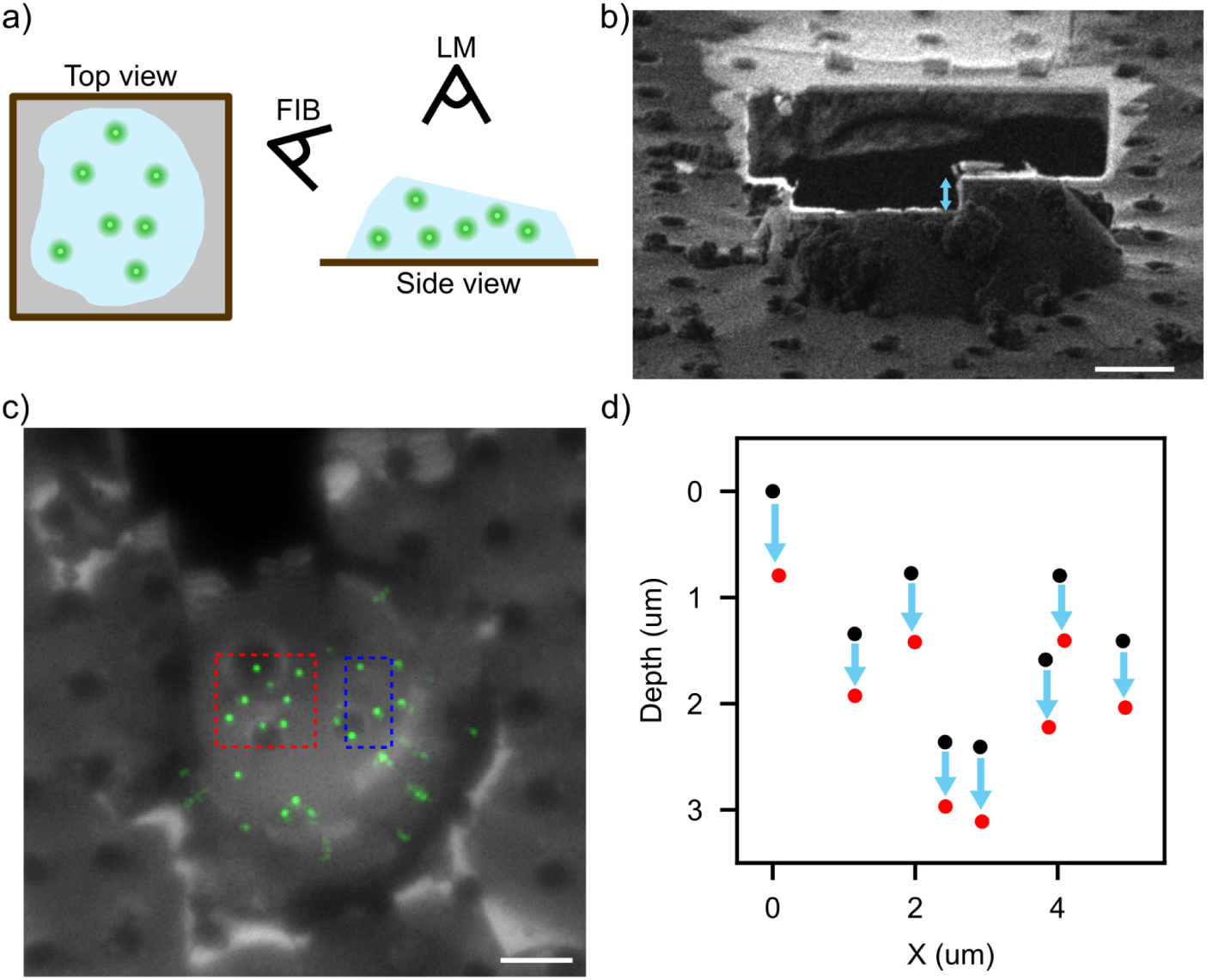
Measurement of focal shift from refractive index mismatch. a) Cartoon of the sample. Green dots represent fluorescent beads suspended in a vitrified water droplet. b) FIB image of a water droplet after the step is cut. Blue arrow highlights material removed between first and second z-stack. c) Z-projected reflection (gray) and fluorescence (green) images of the droplet in (b), taken normal to the grid after the second FIB cut. The blue box shows beads used for registration, and the red box shows beads used for focal shift estimation. Both regions are below the FIB milled surface. d) Black and red dots show estimated position of beads before and after the second FIB cut. Blue arrows indicate the apparent shift downward after material is removed. The average focal shift correction factor from several samples is found to be 1.43. The simulation predicts 1.45 for pure water with a 15° interface. Scale bars 5 μm.

We implemented this depth correction as a workflow in fibsemOS including a precomputed look up table for a wide range of parameters that enables users to apply the depth correction without running simulations. In addition, we implemented automatic fiducial placement into the registration workflow to facilitate more accurate registration between the FIB and the FM images. We next tested and characterized our improved registration workflow on the TFS Arctis FIB SEM instrument.

As a test sample we chose to target centrioles by GFP-labeling the centrosomal protein centrin in immortalized retinal pigment epithelial (RPE-1) cells. We first identify a cell of interest and FIB spot burns are then automatically placed in a pattern over the cell and grid square to be used as milled fiducials (Fig. 4a). An LM z-stack is acquired to determine the rough position of the target within the cell. Based on the rough targeting, stress-relief trenches are milled flanking the target (Wolff et al., 2019), before removing material above the target to create a planar imaging surface. A second fluorescence and reflection z-stack is then acquired normal to the grid (Fig. 4b). Fiducial positions are identified in the reflection channel, while the target is localized in fluorescence (Fig. 4b, inset). Fiducial coordinates are first selected manually and subsequently refined by model-based fitting. An affine transformation with 6 degrees of freedom is then computed to map fluorescence coordinates into the FIB imaging frame. After mapping the apparent fluorescence position onto the FIB image, the depth relative to the milled surface is scaled by the calculated correction factor of 1.5 for these conditions. The predicted target position is then adjusted accordingly prior to start milling around the target.

**Figure 4.**
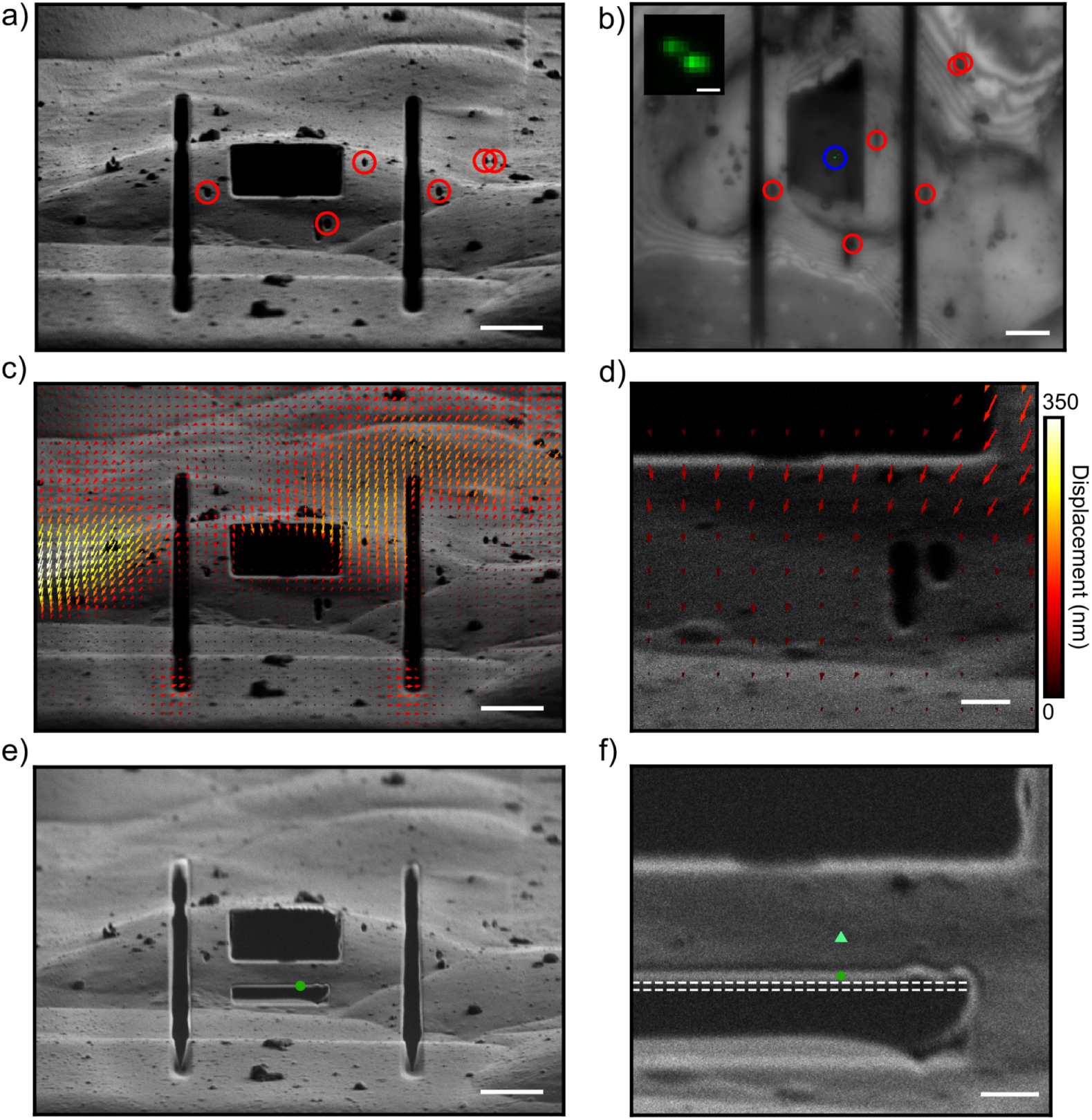
Registration-based targeting. a) FIB image of an RPE-1 cell containing GFP labeled centrioles taken at the milling angle (15°). a) The top surface has been milled away to provide a clean optical window, and stress relief cuts have been made. Red circles mark FIB-milled fiducials used for registration. b) Z-projected reflection (gray) and fluorescence (green) images of the same cell in (a), taken normal to the grid. Red circles highlight the fiducials used for correlation, while a blue circle marks the target. Inset zoomed in on target. c) Deformation field calculated for images before and after real-time fluorescence milling. Arrows show magnitude and direction of deformation for patches of the image, scaled 5x for visibility. d) Zoom in on (c) around the region of interest. e) FIB image taken after real-time fluorescence guided milling. Green dot shows predicted target position. f) Zoom in on (e) around the region of interest. Dark green dot and light green triangle are the predicted target location with and without correction for focal shift. White dashed lines show the beginning and end of the target, determined by live monitoring of fluorescence signal. Scale bars 5 μm (a, b, c, e), 1 μm (d, f), and 500 nm (b inset).

We quantified the targeting accuracy by comparing predicted target positions with the actual object location determined using real-time fluorescence-guided milling (introduced in the next section). Because centrioles are cylindrical structures (250 nm in diameter and 500 nm long), accuracy depends on its orientation relative to the milling direction. In the example shown in Fig. 4f, the predicted position is less than 100 nm from the edge of the object, or within 300 nm from the object’s center. Without focal shift correction, the position is off by almost 1 μm, illustrating the dramatic improvement yielded by incorporating the depth correction into registration workflows. We evaluated the workflow three more times, and found a consistent level of accuracy across all four samples (Fig. S1). While three out of four predicted positions were less than 100 nm from the edge of the object, or less than 300 nm from the object center, in one instance, the object was very close to the surface and the predicted position fell outside of the sample.

Sample deformation during milling remains a limitation. To understand how much sample deformation contributed to the remaining inaccuracy in targeting, we measured sample deformation by comparing images before and after milling. This revealed spatially heterogeneous deformation (Fig. 4c,d), with displacements reaching up to ∼350 nm in some regions. While deformation near the target was smaller (50–100 nm), it remains sufficient to compromise targeting of submicron structures. Increasing the frequency of registration steps can at least partially compensate for this effect, but would reduce throughput without guaranteeing success.

Overall, these results demonstrate that a single-step registration with depth correction enables reliable targeting of objects ≥500 nm in plunge-frozen cells. However, residual errors arising from deformation and uncertainty in sample refractive index limit the reliability of this approach for smaller targets or thicker samples. Achieving higher accuracy with registration alone would require iterative re-registration, incurring substantial time costs. This motivates the use of real-time fluorescence feedback during milling.

### Real-time fluorescence-guided FIB milling

Real-time fluorescence-guided milling provides a direct solution to the limitations of registration-based targeting by coupling material removal to real-time optical feedback. This approach enables precise localization of the target during milling and is particularly effective for submicron structures (Boltje et al., 2022; Wang et al., 2025).

A key requirement is coincident alignment of the FIB and fluorescence microscope, allowing simultaneous access to the same region of the sample. The Thermo Fisher Scientific Arctis PFIB-SEM satisfies this requirement through its integrated geometry, combining an unobstructed TEM-stage with a tilted FIB column, a vertical SEM column, and an inverted fluorescence microscope (Fig. 5a). Despite its potential for real-time fluorescence-guided milling, the standard configuration of this instrument relies on registration-based targeting, precluding this capability.

**Figure 5.**
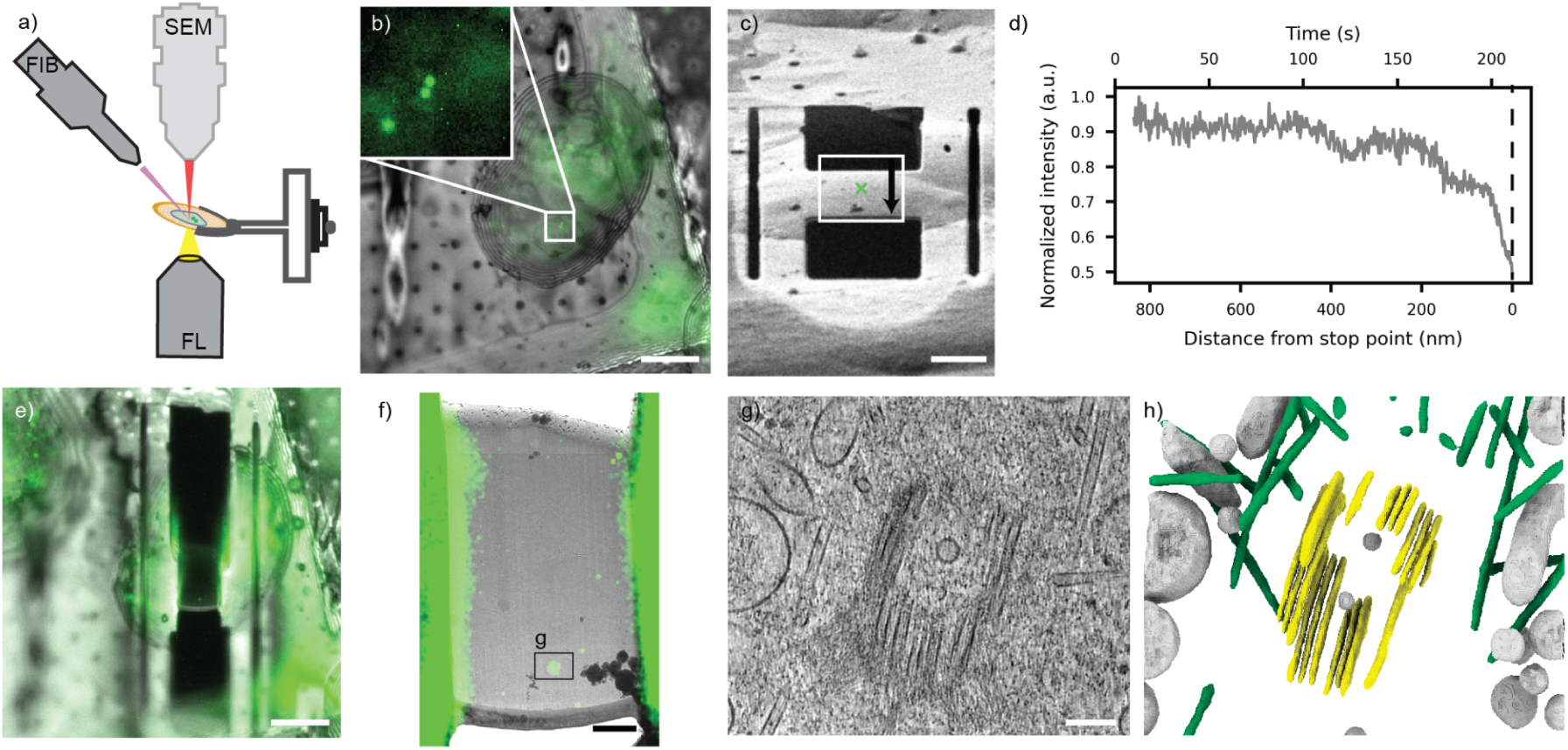
Real-time fluorescence guided FIB milling targeting centrioles. a) Schematic representation of the Arctis sample chamber. The SEM is positioned above the sample, while the integrated fluorescence microscope (FL) is located below the sample. The FIB is oriented at a 52° angle relative to the SEM column. b) Z-projected reflection (gray) and fluorescence (green) images of an RPE-1 cell expressing the centrosomal marker centrin-GFP. Inset: A zoom in on the targeted centrosome as indicated by the presence of the centriole pair. c) FIB image acquired after rough milling. The predicted centrosome target position is indicated by a green x. d) Fluorescence intensity trace recorded during real-time fluorescence-guided milling targeting of the centriole (exposure time 750 ms). e) Reflection image superimposed with the fluorescence image of the final lamella. A bright spot visible in the center of the lamella indicates the presence of the centriole. f) Corresponding overlay of the overview TEM image and the fluorescence image identifying the target location within the lamella. A tilt series was acquired for the highlighted area; a slice through the tomogram confirms the presence of a centriole (g). h) shows a 3D segmentation of the tomogram highlighting membranes (grey), microtubules (green) and centriole microtubule triples in yellow. Scale bars 20 μm (b, c, e), 200 nm (f), 100 nm (g).

To enable this capability, we introduced minimal hardware modifications to allow insertion of the fluorescence objective at the milling angle (−23° stage tilt or 15° milling angle). One change was to remove one of the two cooling braids of the sample holder which eliminated steric interference without compromising cryogenic performance, maintaining temperatures well below the devitrification threshold. Additional software safeguards preventing simultaneous milling and fluorescence imaging were disabled. No measurable degradation of the objective was observed over extended use. SEM imaging was avoided while the fluorescence objective was inserted to prevent electron-beam induced charging of the objective.

Using this modified configuration, we implemented a real-time fluorescence-guided workflow within fibsemOS, built on top of the elements described above. To test and validate the accuracy of this workflow, we turned again to targeting centrioles in centrin-GFP RPE-1 cells. Similar to the registration workflow, a target cell is first identified and fiducial markers are placed to perform registration based rough targeting (Fig 5b) and initial thinning to ∼5 µm (Fig. 5c). Subsequent milling is then performed using the real-time fluorescence approach where fluorescence intensity of the target is recorded while continuously ablating material. For this, the stage is first tilted to the lamella milling angle and the fluorescent signal of the target is brought into focus. A cleaning cross-section pattern (layer-by-layer milling) is then used for milling first from the top and then from the bottom side of the pre-thinned lamella, while the fluorescence intensity within a defined region of interest is continuously monitored. Milling is terminated when a sharp signal drop is observed, corresponding to partial target ablation (Fig 5d, Movie S1). In our examples of labeled centrioles, we specifically targeted one of the two centrioles and stopped the milling after a distinct signal drop (in this case seen after 200 s of milling). The fluorescence image of the final lamella confirms that fluorescence signal remains at the location of the targeted centriole (Fig. 5e,f). Subsequent cryo-TEM imaging confirmed the presence of a centriole in the tomogram acquired at the location indicated by the remaining fluorescence (Fig. 5g). To assess the robustness of the implemented real-time fluorescence guided targeting workflow, we performed 11 experiments and did not observe any failures due to mis-targeting. However, two lamella were lost due to milling failures like lamella breakage or insufficient thinning rather than targeting errors (Fig. S3).

We next applied this workflow to primary cilia, important signaling organelles that have proven difficult to characterize structurally in their native context. Sample preparation has been a key bottleneck; their small diameter (∼200–300 nm) and very low abundance (one per cell) make reliable targeting for FIB milling particularly challenging. Following the workflow described above, we targeted cilia in RPE-1 cells which were fluorescently labeled via heterologous expression of a halo-tagged serotonin receptor 6 (5-HTR6) (Deo et al., 2019). Essentially, we monitored fluorescence during milling to decide when to stop or adjust the milling (Fig. 6, Movie S2). Cryo-TEM imaging confirmed successful capture of ciliary cross-sections (Fig. 6d,e,f), and tomograms revealed the expected membrane and microtubule organization characteristic of primary cilia (Kiesel et al., 2020) (Fig. 6f, Fig. S4). Across 5 additional experiments targeting primary cilia, we again did not observe any targeting errors and the lone failure was due to lamella instability.

**Figure 6:**
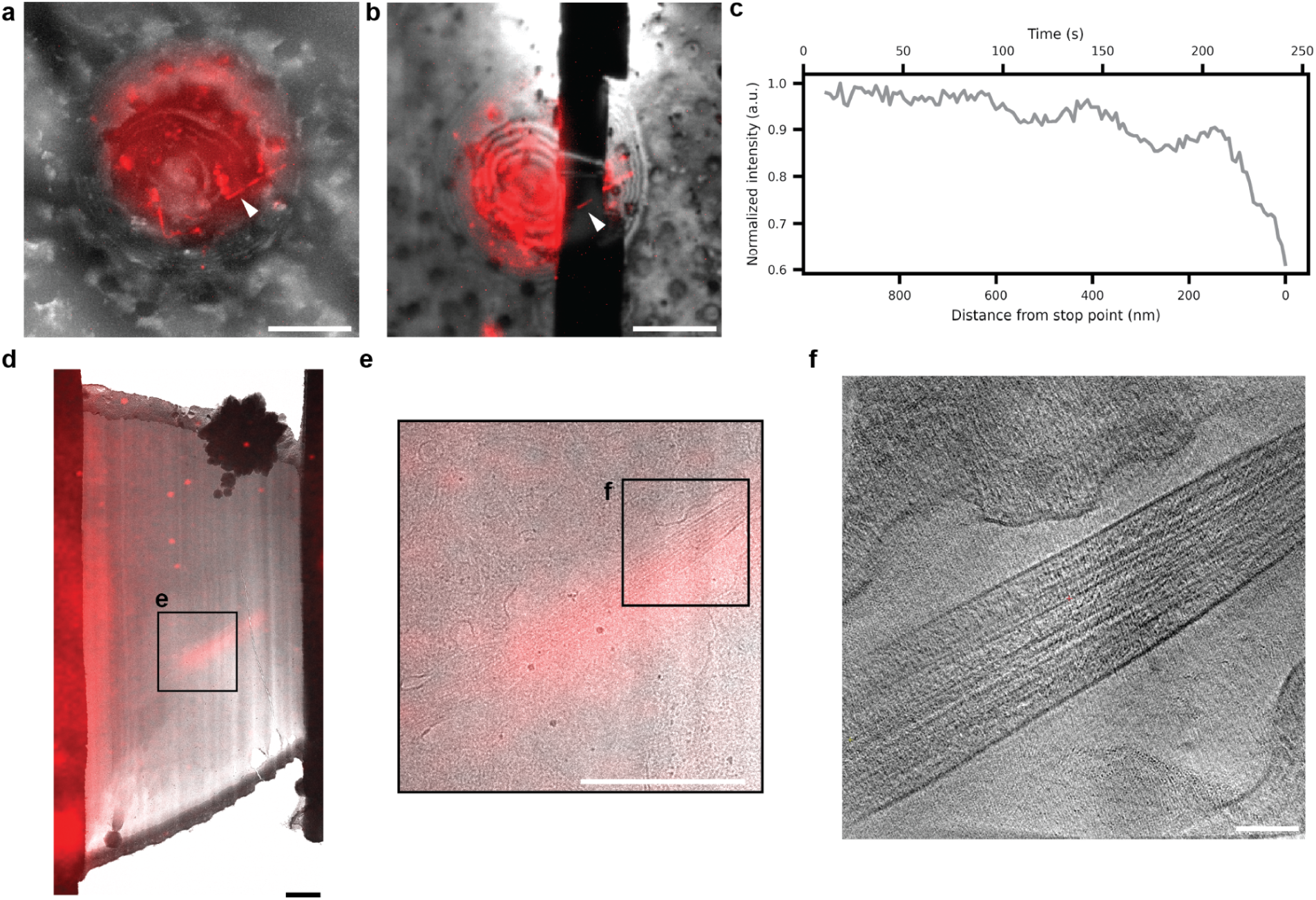
Real-time fluorescence-guided FIB milling enables capture of primary cilia for cryo-ET. a) Cryo-fluorescence image overlaid on a reflection image of RPE-1 cells with labeled primary cilia shown in red. The cilia-specific 5-HTR6 receptor is labeled with a JF552-HaloTag. The targeted cilium is indicated by a white arrowhead. b) Same imaging modality and sample position as in a) following real-time fluorescence-guided milling, confirming retention of the targeted cilium within the prepared lamella. c) Real-time fluorescence intensity as a function of time, illustrating the progressive decrease in signal as cellular material is removed during milling. d) TEM overview of the lamella overlaid with the final cryo-fluorescence image, indicating the position of the cilium within the lamella. The cilium is clearly identifiable in the TEM image, as shown at higher magnification in e). f) Tomographic slice through the cilium at the position indicated in e). Scale bars 10 µm (a, b), 1 µm (d, e), 100 nm (f).

The fluorescence trace in Fig. 6c suggests an interference pattern as previously described (Sica et al., 2026), an effect we also observed in experiments on fluorescent beads in water droplets (Fig. S5). Sica et al. used these interference patterns to accurately estimate the distance from the object of interest during milling to target even smaller structures (<200 nm). We identified such interference patterns in a subset of our data when there was high signal over background, suggesting that despite differences between the experiments performed by Sica *et al* and this work, the interference effect can be used to implement more accurate targeting on the Arctis in the future.

Together, these results demonstrate that real-time fluorescence-guided milling on a commercially available platform enables robust and efficient targeting of low abundance, sub-500 nm structures, overcoming a central limitation of in situ cryo-ET.

## Discussion

Here, we present two improved workflows for fluorescence-guided targeting during cryo-FIB milling. Our approaches address complementary targeting regimes. Registration-based correlation, which is compatible with any cryo-fluorescence microscope (integrated or standalone), provides reliable single step targeting of objects ≥500 nm. Real-time fluorescence-guided milling, which requires an LM-FIB coincidence point, is critical for smaller structures and overall the more reliable approach if an appropriate instrument is available. Together, with these two approaches we establish a practical framework for targeting cellular structures across a broad range of sizes and abundances.

Our registration-based workflow integrates several advances from across the field into a streamlined and user-accessible pipeline implemented in fibsemOS. By incorporating automatically generated FIB-milled fiducials, we provide robust and reproducible landmarks independent of intrinsic sample features. In combination with coordinate transformation approaches inspired by 3DCT (Arnold et al., 2016; Klumpe et al., 2021), this enables automated mapping of fluorescence coordinates onto FIB images and precise placement of milling patterns. The integration of these different components into a single, coherent workflow substantially lowers the barrier to adoption and improves experimental throughput and success rates. Crucially, this workflow does not depend on any specific instrument geometry and can be performed on any FIB SEM instrument in combination with a cryo light microscope.

Beyond integration, a central contribution of this work is the introduction of a physics-based correction for focal shift arising from refractive index mismatch between the sample and vacuum. This effect can displace apparent target positions by microns, sufficient to place targets entirely outside the final lamella, yet is often neglected or only approximately corrected. Using vectorial diffraction simulations validated by experimental measurements in vitreous ice, we establish a systematic framework for determining the appropriate depth scaling factor under defined imaging conditions (tilt, refractive index, NA, etc). By computing this factor across a broad parameter space, we provide a resource that can be directly applied across instruments and sample geometries without requiring additional modeling.

With these improvements, we show that single-step registration enables reliable targeting of objects on the order of ∼500 nm or larger in plunge-frozen samples. Sample deformation during milling emerges as the major residual source of error. In addition, the scaling factor is linearly dependent on the refractive index of the sample and variations in refractive index remains a source of uncertainty (Fig. S2). The refractive index of vitrified cellular material has not been well measured, and refractive index is known to vary throughout the cell at room temperature (Baczewska et al., 2021; Schürmann et al., 2016). Additionally, the model does not account for sample or microscope-specific optical aberrations.

This single-shot targetable regime of up to 500 nm size objects for our improved registration based workflow encompasses many biologically relevant structures and provides a practical solution for experiments where instrument geometry precludes simultaneous fluorescence imaging during milling. Further, registration remains an essential component of our accurate real-time fluorescence-guided workflows, providing the initial localization that is subsequently refined. To target structures smaller than 500 nm, iterative registration can partially mitigate targeting inaccuracies from sample deformation but it does not guarantee successful targeting of smaller objects, motivating the need for real-time fluorescence targeting approaches.

To address this limitation, we implement real-time fluorescence-guided FIB milling on a commercially available Thermo Fisher Scientific Arctis platform. We demonstrate that this workflow achieves targeting accuracies of greater than 90%, comparable to previously reported implementations on custom build instruments (Boltje et al., 2022; S. Li et al., 2023; Wang et al., 2025). Importantly our implementation on the Arctis only requires a minimal hardware modification. Specifically, a straightforward adjustment to the sample holder enables simultaneous access of the fluorescence objective at the milling angle, providing a simple and accessible upgrade path. Although the current implementation requires manual intervention, several steps—including coarse correlation, initial milling, and identification of the fluorescence drop-point—are amenable to automation, which is currently underway. Integration of these elements on high-throughput milling platforms such as the Arctis has the potential to enable scalable acquisition of tomography data on fluorescently labeled targets, a critical requirement for in situ structural biology.

We validate our implementation of a real-time fluorescence guided FIB milling workflow using individual centrioles and primary cilia, demonstrating reliable targeting of small, rare, and biologically significant organelles. By enabling precise and reproducible targeting, the approaches described here open the door to systematic in situ structural studies of these and other low-abundance cellular features. More broadly, these methods are well suited to applications requiring selective targeting of specific instances of otherwise abundant structures, such as studies combining cryo-ET with fluorescence biosensors.

Finally, the semi-destructive nature of real-time fluorescence-guided milling imposes a practical lower bound on target size, estimated here at ∼150 nm. For smaller objects, alternative strategies such as interferometric targeting may be required. Notably, our observations suggest that the interferometric signal previously reported for gallium FIB systems (Sica et al., 2026) is also present under plasma FIB conditions and with a different stage geometry, although notably with a different phase. In our observations, the object is destroyed at an interference maximum, suggesting the phase is not inverted upon reflection at the interface, consistent with a reflection from high to low refractive index. Sica et al. observed the opposite, which was attributed to a gallium implantation or redeposition layer creating a reflection from low to high index. For the plasma FIB used in this work, that gallium layer does not exist. At the same time, observation of interference is complicated by stray fluorescence from the lamella itself. Because of total internal reflection, fluorescence excited throughout the cell or droplet is guided along the lamella and concentrated at its edges. This edge fluorescence can overlap spatially with the target signal, obscuring the interference fringes. The effect is more pronounced on the Arctis than on the system used by Sica et al., because the steeper angle between the lamella and optical axis increases the overlap between the bright edge and the region of interest, which might explain why we only observed the interference in a subset of our data acquired on the Arctis. Further characterization of this signal and its potential for high-precision targeting represents an important avenue for future work, especially for fluorescence-guided FIB milling and the targeting of sub-150 nm structures. Nonetheless, the advances presented here already substantially expand the range of targets accessible to cryo-ET, including many that were previously out of reach.

## Methods

### Cell culture and grid preparation

hTERT RPE-1 cells stably expressing 3×GFP-centrin (centrin–GFP) were obtained from the Cheeseman laboratory (Whitehead Institute, MIT) and maintained as previously described (Su et al., 2016).

Quantifoil Au 200 grids (SiO₂ R1/4) were glow discharged for 30 s (Pelco EasyGlow) and UV-sterilized for ∼20 min. Individual grids were placed in 35-mm glass-bottom dishes (No. 1.5, uncoated; MatTek) on 2 µl PBS droplets, with surface tension maintaining grid position.

Cells were trypsinized, counted, and resuspended at 3 × 10⁵ cells ml⁻¹ in complete medium. Three milliliters of cell suspension was added per dish with minimal disturbance to grids. Dishes were placed in secondary containment and incubated at 37 °C and 5% CO₂ for ∼4 h.

### Cell culture and grid preparation for primary cilia cryo-ET

Human hTERT RPE-1 cells (ATCC CRL-4000) were used for primary cilia imaging. These cells were maintained in DMEM/F-12 (1:1) supplemented with tetracycline-free 10% fetal bovine serum (FBS) at 37°C in a humidified atmosphere with 5% CO₂. To label primary cilia, a modified RPE-1 cell line stably expressing a tetracycline-inducible HTR6-HaloTag fusion protein was used, as previously described (Deo et al, 2019; Sheu et al., *Cell* 2022). At approximately 80% confluence, the growth medium was replaced with serum-free DMEM/F-12 supplemented with 100 ng/ml doxycycline to simultaneously induce HTR6-HaloTag expression and promote primary cilia formation through serum starvation. Cells were maintained under these conditions for 48 hours.

To fluorescently label cilia, cells were incubated with 250 nM Janelia Fluor 552 HaloTag ligand (JF552-HTL) in serum-free media for 30 min at 37°C. Following labeling, cells were washed twice with phosphate-buffered saline (PBS) to remove unbound dye. Cells were then detached from the culture dish using a cell scraper and resuspended in serum-free DMEM/F-12. Cell suspensions were applied onto freshly prepared EM grids at 50,000-100,000 cells per grid.

Before cell seeding, grids were micropatterned using the PRIMO micropatterning device (Alveole). EM grids (200 mesh gold Quantifoil R2/1 with SiO₂ support film) were first glow-discharged to render the surface hydrophilic (Pelco easy glow). Grids were then incubated with poly-L-lysine-*graft*-polyethylene glycol (PLL-*g*-PEG; SuSoS) as an antifouling passivation layer. Circular adhesion zones of 60 µm diameter were defined at the center of each grid square using the PRIMO maskless photopatterning system (Alvéole) with the PLPP photoinitiator and the Leonardo software, which automatically aligns patterns to the grid mesh geometry (Engel et al., 2019). UV exposure locally degraded the PLL-*g*-PEG to promote cell adhesion within those confined areas. Cells were seeded onto patterned grids and allowed to adhere overnight at 37°C prior to plunge freezing.

### Grid vitrification

RPE-1 cell–seeded grids were vitrified using a Leica GP2 plunge freezer. The chamber was maintained at 37 °C and 80–90% relative humidity. Grids were blotted for 1–10 s and plunge-frozen in liquid ethane at −180 °C. All subsequent handling steps were performed under liquid nitrogen. Grids were clipped and stored in liquid nitrogen until further use.

### Bead droplet sample prep

Bead samples were produced using a gas dynamic virtual nozzle (GDVN) microfluidic aerosol system which has been previously described (Yoniles et al., 2024; Sica et al., 2026). Briefly, the GDVN was placed ∼ 7 cm away from the plunging path of the grid to allow for the grid to pass through the aerosolized bead samples. A flow rate of 0.12 ml/min was used along with intentionally non-glow discharged copper Quantifoil grids (EMS Q2100CR2) to produce high contact angle droplets approximately 10-20 μm in diameter. The two different bead samples used were a 0.01% solids aqueous sample containing 200 nm orange (540/560 nm) polystyrene beads (FluoSpheresTM F8809) and a 0.005% solids aqueous sample containing 100 nm blue (350/440 nm) polystyrene beads (FluoSpheresTM F8797).

### Registration targeting of centrioles in RPE-1 cells

Grids with RPE-1 cells expressing GFP labeled centrin were prepared as described above and loaded onto a Thermo Fisher Arctis PFIB-SEM equipped with an integrated fluorescence light microscope (iFLM) with a 100x 0.75 NA objective. Grids were coated with organoplatinum material using the gas injection system to protect the leading edge during milling, and then sputter coated with platinum for surface conductivity. The instrument was controlled using the fibsemOS software package described in more detail below. A FIB tileset was acquired at the milling angle of 15° to identify cells in good position for milling. A pattern of FIB spot burns was automatically placed over the cell to be used as fiducials with a beam current of 100 pA and exposure times of 1 to 10 s. LM z-stacks were acquired at a stage position of -180° (optical axis normal to the grid, looking at the top of the sample). The milled fiducials were used to roughly correlate the LM and FIB images so the top trench could be cut above the centrioles, with stress relief cuts to each side. After these cuts, a high resolution FIB image was acquired, followed by an LM z-stack with 250 nm steps and fluorescence and reflection color channels. These images were used to predict the target locations. Five or more spot burns were selected manually in FIB and reflection images. In the FIB images, positions were refined by fitting a 2D gaussian with negative amplitude to the holes. For reflected light images, a 1D gaussian was fit to the axial profile to find the focal position, and a 2D gaussian was fit to an image interpolated to that plane. The same procedure was applied to the target in the fluorescence image. When this fitting routine failed, manually selected positions were used instead. To account for focal shift, the top surface of the cut face was manually selected in the FIB image, and the depth of the predicted target position relative to that surface was multiplied by the calculated scaling factor.

To check the accuracy of this prediction, real-time fluorescence-guided milling was done following the procedure outlined below. With the stage at the milling angle, fluorescence signal from the object was monitored while the sample was milled from below. When the signal began to drop, the milling was paused and a FIB image was taken. Then milling was resumed until the object had been milled away, and another FIB image was taken. These images were registered to the FIB image used for registration by SIFT registration in Fiji (Lowe, 2004; Schindelin et al., 2012) calculated for a portion of the image which should not experience deformation, such as a grid bar. The distance of the predicted target position from the bottom cut face is the targeting error, and was found to be on the order of 100-300 nm.

### Bead focal shift measurement

Grids with bead droplets were prepared as described above and loaded onto a Thermo Fisher Hydra PFIB-SEM with a Delmic Meteor equipped with a 50x 0.8 NA objective. Grids were coated with organoplatinum material using the gas injection system to protect the leading edge during milling, and then sputter coated with platinum for surface conductivity. The stage was tilted for a milling angle of 15°, and the top of the droplet was removed for a smooth, known surface for optical imaging. The stage was moved to the LM position, and z-stacks were acquired top down normal to the grid with 250 nm steps. The stage was returned to the milling position, and 1 to 3 μm of material was removed from one side of the droplet. Z-stacks were acquired again with the same parameters.

### Real-time fluorescence-guided targeting

Grids with RPE-1 cells containing GFP labeled centrin or JF552 labeled 5HT6-halotag were prepared as described above and loaded onto a Thermo Fisher Arctis. The samples were coated with organometallic platinum with 120 s GIS and a subsequent 120 s platinum sputter coating. To prevent excessive reflection when imaging through the bottom of the sample, it is important to omit the initial platinum sputter coat. Cells of interest are identified in FIB tilesets taken at the milling angle, and positioned at the FIB-SEM coincident point. Then rough registration targeting is carried out as described above to place rough milling patterns around the target. No depth correction was applied here, because the object only needs to be captured between two rough cuts, whose spacing was set to 5 μm, and the total sample thickness is only ∼ 10 μm. After rough cuts, the iFLM is inserted and focused on the fluorescent target with the stage at the milling angle. This is only possible because of the modified sample holder. A region of interest is defined in the coincidence milling software tool integrated in autolamella (fibsemOS). During milling, the intensity within this window is averaged and monitored to determine when to stop the milling. A milling current of 100 pA (at 30 kV) was used during the real-time fluorescence-guided milling. Depending on the width of the cleaning-cross-section pattern, this corresponds to a milling speed of 6 to 4 nm/s. Real-time fluorescence-guided milling was performed both from top-down and bottom-up milling direction using a cleaning-cross section milling pattern. Finally the lamellae were polished using a milling current of 30 pA.

### FIBSEMOS

As part of this work, several new features have been implemented in fibsemOS for the AutoLamella application. The registration based workflow from 3DCT (Arnold et al., 2016; Klumpe et al., 2021) is integrated directly into the Graphical User Interface (GUI). This includes the standard workflow as well as improvements discussed in this work; refractive index correction, a correction factor calculator, and metadata parsing for Delmic Meteor images. The spot burn fiducial preparation is included as a standard, executable task within the workflow. A dedicated GUI allows users to precisely select spot positions and define relevant parameters (beam current, dwell time). Both the spot burn fiducials and the registration workflow are available on all fibsemOS supported microscopes. Specific to the TFS Arctis microscope, we have added fluorescence microscope control. This control supports multi-channel z-stack acquisition, auto-focus, overview acquisition, and full metadata management. The real-time fluorescence-guided milling widget described in this work is integrated with the AutoLamella workflow. This widget allows the user to select the region of interest and monitor fluorescence intensity in real-time during the milling process. The software will be released publicly under an open-source license at https://github.com/fibsem-os/fibsem-os.

### CryoET

Milled samples were immediately transferred into a TFS Krios G4 equipped with a Falcon4 direct detector and a Selectris X energy filter. Images were acquired using the TOMO5 software. Lamella were identified in overview images and Search maps were collected at a magnification of 6500. For Search map and tilt series a pre-tilt of -15° was set to compensate for the milling angle of 15°. Tilt series were acquired with a pixel size of 1.95 Å/px. The targets were manually identified from the search maps, and tilt series were acquired from -60° to 30°, starting from -15°, with 3° tilt increments using a dose symmetric tilt scheme and a total dose of 120 e/Å^2^. Tilt series were motioncorrected and automatically reconstructed into tomograms using Aretomo3 2.2.9 (Peck et al., 2025). Tomogram slices were visualized in 3dmod 5.1.3 (Kremer et al., 1996). Tomogram segmentations were performed using Copick, a data-centric API that streamlines model training and inference workflows (Ermel et al., 2026). Membrane segmentations were generated using Membrain-seg (Lamm et al., 2024) in combination with copick-torch, while microtubule and cilia segmentations were obtained using nnInteractive (Isensee et al., 2025) via the napari-copick interface. Segmentations were visualized in UCSF ChimeraX 1.10 (Meng et al., 2023). Post-milling fluorescence images were overlaid to TEM overview images by manual landmark-based affine registration of the FM image into the TEM coordinate frame.

## Supporting information

Optics simulation code

## Acknowledgments

We thank Thermo Fisher Scientific for collaborating on this project, in particular for their support in providing the modified sample holder for the Arctis, productive discussions and technical assistance. We thank Liz Montabana, Misha Kopylov, Hrishita Sitwala for assistance in TEM data collection, Nikki Jean for help screening samples and Utz H. Ermel for assistance with tomogram segmentations.

D.P., S.B., C.V., J.S., D.A.A, B.C., J.P., G.G. are internally funded. Funding was provided to P.C. and S.K. from the Chan Zuckerberg Initiative, grant number 2025-366327. Research in the lab of S.K. is supported by the Austrian Academy of Sciences and Boehringer Ingelheim. C.A. and P.D.D. were supported by the Laboratory Directed Research and Development program at SLAC National Accelerator Laboratory, under contract DE-AC02-76SF00515.

We also thank the Cheeseman lab (Whitehead Institute for Biomedical Research and the Department of Biology at MIT, Cambridge, MA, 02142 for the hTERT RPE-1 - cells expressing 3xGFP-centrin.

