## Supplementary figures and notes for "High accuracy fluorescence-guided cryo-focused ion beam milling"

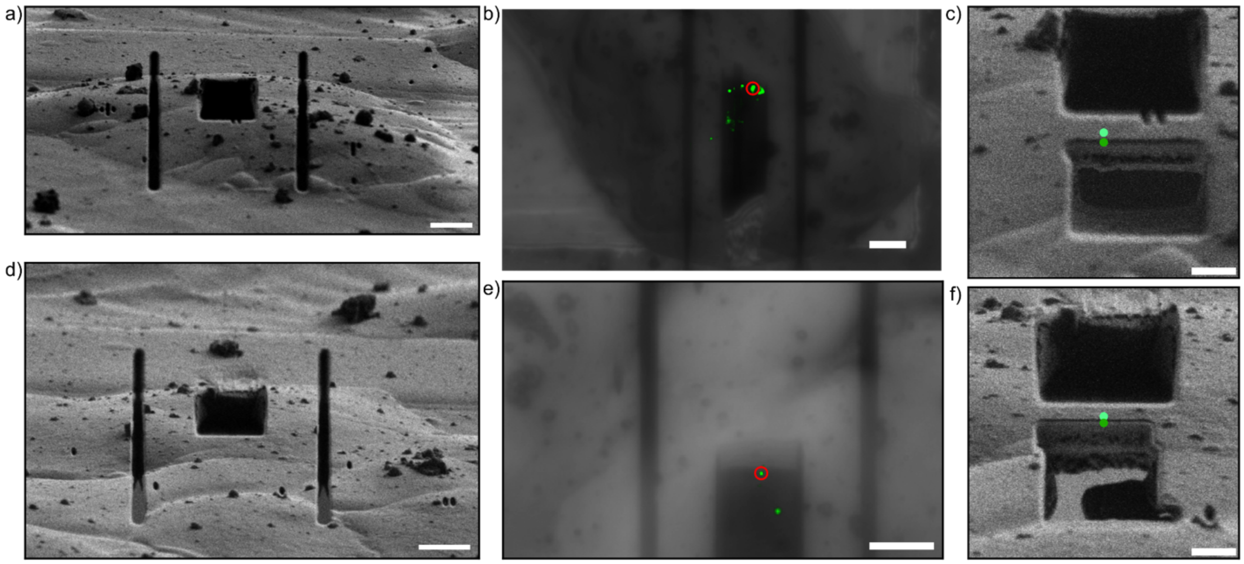


Figure S1. Repeated registration accuracy testing. a,d) FIB images after site preparation. b,e) LM z-projections showing fluorescent target. c,f) Predicted target position with (dark green) and without (light green) focal shift correction displayed on FIB image. The bottom cut was performed while watching fluorescence signal, and milling was stopped after the target was completely milled through. In both cases, the predicted target position (with focal shift correction applied) lies right under the cut face, suggesting good agreement with the actual object position. Scale bars 5 um (a, b, d, e) and 2 um (c, f).

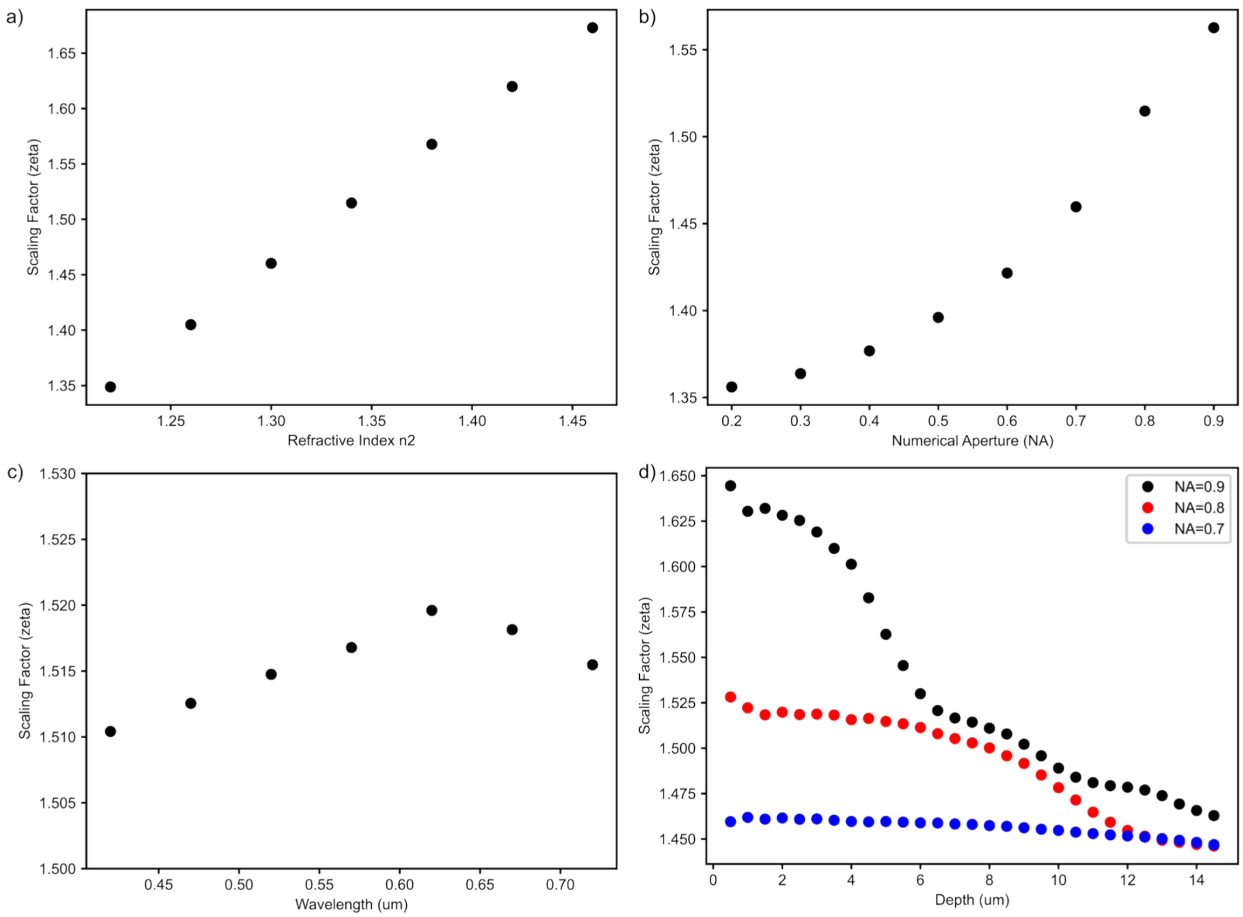


Figure S2. Computed scaling factors vs: (a) refractive index (n2) with tilt = 15°, NA = 0.8, wavelength = 520 nm, object depth = 5 um; (b) numerical aperture (NA) with tilt = 15°, n2 = 1.34, wavelength = 520 nm, object depth = 5 um; (c) wavelength with tilt = 15°, n2 = 1.34, NA = 0.8, object depth = 5 um; and (d) depth with tilt = 15°, n2 = 1.34, wavelength = 520, and NA = 0.7, 0.8, and 0.9.

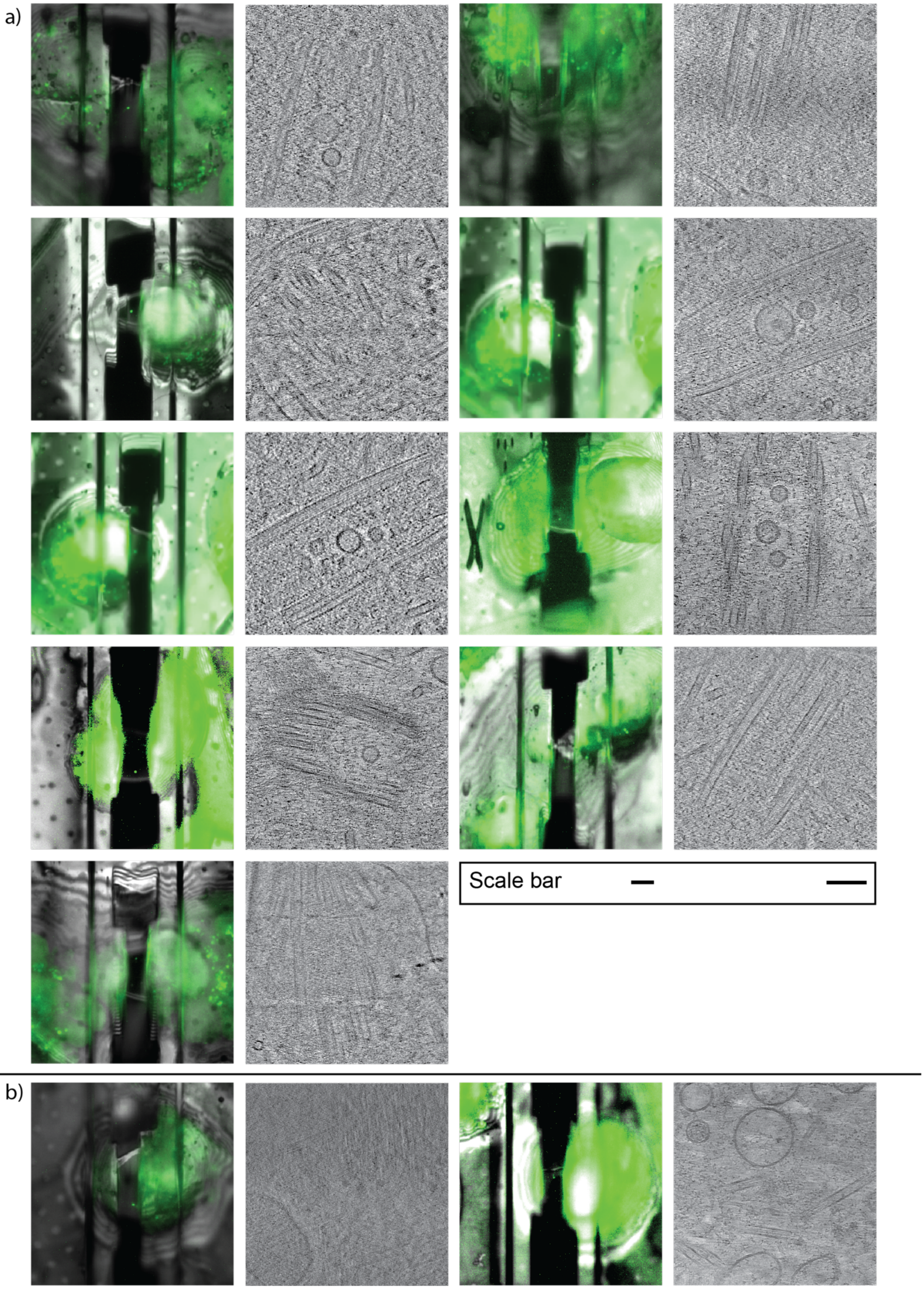
a)
Scale bar
b)

Figure S3. Overview of the real-time fluorescence guided milling experiments targeting centrosomes. The overlay of reflection and fluorescence images (scale bar 5 μm) of the final lamellae are displayed together with a slice of the respective tomogram (scale bar 100 nm). a) 9 centrioles could be successfully targeted and imaged. b) In two cases the fluorescence signal is still visible, but the TEM lamella preparation failed.

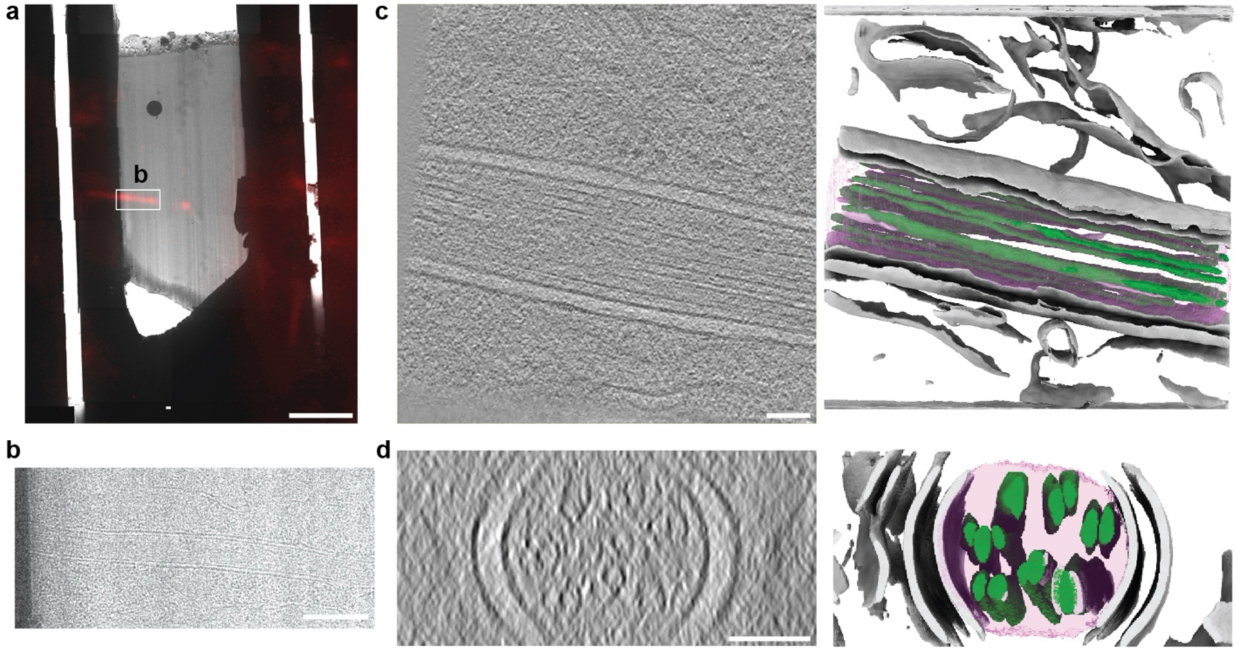


Figure S4: Additional example of a successfully targeted cilium through real-time fluorescence guided FIB milling. a) Overlay of the final fluorescence image of the lamellae and the TEM overview maps of the lamellae indicating that a section of the targeted cilium remained. Scale bar: 5 um. b) Zoom in into the lamella overview shown in a, showing the cilium section captured. Scale bar: 500 nm c) XY-slice through a tomogram acquired along this cilium and the corresponding segmentation on the right. Scale bar: 100 nm d) Zoomed in slice through the tomogram shown in b along the direction of the cilium, showing a cross section of the captured cilium and the corresponding segmentation on the right. Scale bar: 500 nm. In the segmentations grey surfaces are membrane, the pink volume is the cilium, green tubules are axonemal microtubules and microtubule doublets.

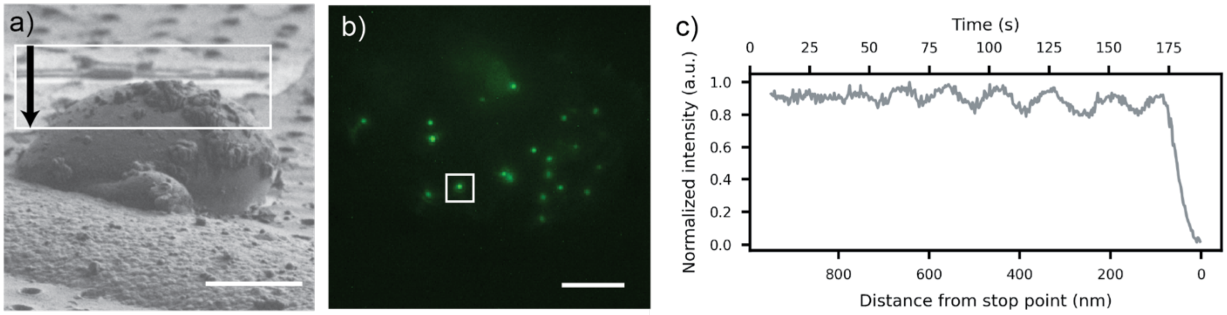


Figure S5. Overview of the real-time fluorescence-guided milling experiment targeting fluorescent beads (excitation wavelength 365 nm, diameter 100 nm) in water droplets. a) FIB image of a water droplet containing fluorescent beads prior to milling (scale bar 10 µm). b) Corresponding fluorescence image showing the spatial distribution of beads within the droplet (scale bar 5 µm). c) Fluorescence intensity trace recorded during the milling experiment for the bead highlighted by the white square in b). Milling proceeds from the top of the droplet toward the support film. Interference fringes are visible in the signal. A two-dimensional Gaussian fit was applied to correct for background reflections from the surroundings.

### Supplementary note 1 – optics simulation

Problem statement:

Given experimentally defined parameters (interface tilt, wavelength, NA, n2) and the object's depth relative to the interface, estimate the object's apparent position.

Defining the coordinate system:

For this calculation, we work in a right-handed laboratory coordinate system $(x, y, z)$ with the optical axis of the microscope along $\hat{z}$. The objective lens collects light propagating in the $+z$ direction.

A planar interface passes through the origin with normal $\hat{n} = (\sin\tau, 0, \cos\tau)$, where $\tau$ is the tilt angle about the y axis. The normal points form the sample medium (refractive index $n_2$) toward the vacuum (refractive index $n_1 = 1$).

The object is modeled as a point emitter at $r_o = (0, 0, -z_O)$ in medium $n_2$.

Image calculation:

For a given object position and set of experimental parameters, we will calculate the point spread function (PSF) with vectorial diffraction theory. We assume the object is an isotropic point emitter, and the optical system is aberration-free. To calculate the PSF, we start by constructing the pupil function.

While we are interested in the emission direction, by optical reversibility this is equivalent to the focusing direction which we will use to calculate the pupil function. We start by constructing a k-space grid in the back focal plane of the objective, limited by the numerical aperture. The Richards-Wolf integral is used to map these k vectors in the back focal plane to vectors on a curved reference sphere. Specifically for a pupil point $(k_x, k_y)$ in medium $n_1$, the propagation direction on the reference sphere is:

$$s_1 = \frac{1}{n_1 k_0}\left(k_x, k_y, k_{1z}\right), \text{ where } k_{1z} = \sqrt{(n_1 k_0)^2 - k_x^2 - k_y^2}$$

where $k_0 = 2\pi/\lambda$ is the vacuum wavenumber. The polar angle $\theta_1$ satisfies

$\cos\theta_1 = k_{1z}/(n_1 k_0)$, and the azimuthal angle is $\phi = \arctan(k_y/k_x)$. The numerical aperture restricts the pupil to $\sqrt{k_x^2 + k_y^2} \leq k_0 NA$. The Jacobian of this mapping introduces the apodization factor $A(\theta_1) = \frac{1}{\sqrt{\cos\theta_1}}$ to satisfy the Abbe sine condition.

Snell's law is then applied refract the rays from vacuum to sample medium at the interface. We use the vectorial form of Snell's law:

$$s_2 = \frac{n_1}{n_2}(s_1 - (s_1 \cdot \hat{n})\hat{n}) + \sqrt{1 - \frac{n_1^2}{n_2^2}(1 - (s_1 \cdot \hat{n})^2)}\ \hat{n}$$

Rays found to undergo total internal reflection (TIR) are excluded from further calculation.

To map polarization from k-space, we use the equations:

$$E_{inc}^{(x)}(\theta_1, \phi) = \begin{pmatrix} \cos^2\phi \cos\theta_1 + \sin^2\phi \\ \sin\phi\cos\phi(\cos\theta_1 - 1) \\ -\cos\phi\sin\theta_1 \end{pmatrix}$$

$$E_{inc}^{(y)}(\theta_1, \phi) = \begin{pmatrix} \sin\phi\cos\phi(\cos\theta_1 - 1) \\ \sin^2\phi\cos\theta_1 + \cos^2\phi \\ -\sin\phi\sin\theta_1 \end{pmatrix}$$

To calculate Fresnel transmission at the interface, polarization is decomposed into s and p components relative to the plane of incidence at the interface. The plane of incidence is spanned by $\widehat{s_1}$ and $\hat{n}$ with local basis vectors $\widehat{e_s} = \frac{\widehat{s_1} \times \hat{n}}{|\widehat{s_1} \times \hat{n}|}$ and $\hat{e}_{p_1} = \widehat{e_s} \times \widehat{s_1}$. Therefore, for x,

$E_s^{(x)} = E_{inc}^{(x)} \cdot \widehat{e_s}, E_p^{(x)} = E_{inc}^{(x)} \cdot \hat{e}_{p_1}$.

Fresnel coefficients ($t_s$ and $t_p$) are calculated for propagation from $n_2$ to $n_1$, since we care about the emission direction. The, the transmitted field is reconstructed as:

$$E_{trans}^{(x)} = t_s E_s^{(x)} \widehat{e_s} + t_p E_{p_1}^{(x)} \widehat{e_{p_2}}$$

Next, we calculate the phase accumulated between the source and the planar interface. In the source medium 2, every wavevector $k_2 = (k_{2x}, k_{2y}, k_{2z})$ has spatial dependence $e^{ik_2 \cdot (r - r_0)} = e^{ik_2 \cdot r} \cdot e^{-ik_2 \cdot r_0}$. From Snell's law, we have $k_1 = k_2 + \Delta k \hat{n}$.

In medium 1, the electric field is $E_1(r) \propto e^{ik_1 \cdot r + i\phi}$. The boundary condition at the interface requires $e^{ik_1 \cdot r_{int} + i\phi} = e^{ik_2 \cdot (r_{int} - r_0)}$.

For any point $r_{int}$ on the interface, $\hat{n} \cdot r_{int} = 0$ since the interface passes through the origin, so $k_1 \cdot r_{int} = (k_2 + \Delta k \hat{n}) \cdot r_{int} = k_2 \cdot r_{int}$. Substitution into the above equation gives:

$$\phi = -k_2 \cdot r_0 = k_{2z} z_O = k_0 n_2 s_{2z} z_O$$

Combining these results gives the vectorial pupil function for input polarization $\hat{e}_{in} \in \{\hat{x}, \hat{y}\}$ as:

$$P^{(in)}(k_x, k_y) = \frac{1}{\sqrt{\cos\theta_1}} e^{ik_{2z} z_O} E_{trans}^{(in)}(k_x, k_y)$$

Where $E_{trans}^{(in)}(k_x, k_y)$ is the transmitted filed defined above. The three Cartesian components of each pupil function are stored as separate 2D arrays:

$P_{0,j}^{(in)}(k_x, k_y) = P^{(in)} \cdot \hat{e}_j$, for $j \in \{x, y, z\}$ and input polarization $(in)$.

To calculate a given real-space image, this pupil function is combined with a propagator $H(k_x, k_y; z_N) = e^{-ik_{1z}z_N}$ for nominal focal depth $z_N$. The electric field is therefore:

$$E_j^{(in)}(x, y, z_N) = \mathcal{F}^{-1}\left\{P_{0,j}^{(in)}(k_x, k_y) \cdot H(k_x, k_y; z_N)\right\}$$

For unpolarized fluorescence emission, the intensity is the sum over the two orthogonal input polarizations:

$$I(x, y, z_N) = \sum_{in \in \{x,y\}} \sum_{j \in \{x,y,z)} \left|E_j^{(in)}(x, y, z_N)\right|^2$$

Apparent position determination:

The above calculation is used to create a series of 151 images spanning 10 um around the expected apparent position based on the paraxial approximation $z_A = z_O * \frac{n_1}{n_2}$ for apparent depth $z_A$ and object depth $z_O$. The images are calculated with a pixel size of 40 nm. The max pixel value is taken from each of these images to create a z profile. This metric was chosen for its insensitivity to the asymmetric shapes of the highly aberrated PSFs produced at high tilt or large depth at high NA. The max of this z profile is used as a course estimate of the in-focus position. This estimate is refined by a local (11 point wide) O(2) polynomial fit. Again, this metric was chosen to be minimally sensitive to the highly skewed axial profiles produced in the more extreme conditions. Choosing an alternative metric, such as the centroid, best-fit gaussian, etc., significantly effects the calculated apparent position for these extreme cases, but has a minimal effect at more modest tilts and depths most relevant to this experiment. The depth scaling factor is calculated as $\zeta = z_O / z_A$.

Claude and Gemini were used to assist with these calculations.

Bead focal shift measurement

To relate the material removed measured in FIB view to material removed along the optical axis:

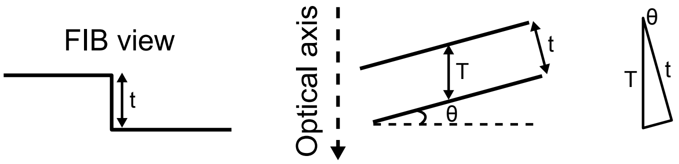


$$T = t/\cos\theta$$

Where T is the material removed along the optical axis, t is apparent material removed from the FIB view, and $\theta$ is the milling angle.

To calculate correction factor $\zeta$ from the bead focal shift measurements:

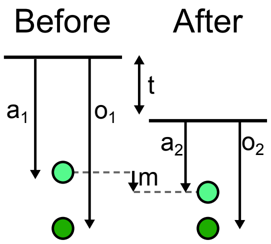


t is the material removed on the optical axis, $a_1$ is the apparent depth after the first cut for object depth $o_1$, $a_2$ is the apparent depth after the second cut for object depth $o_2$, and m is the measured change in apparent depth. To calculate $\zeta$ given t and m, we use the known relations

$$o_1 = \zeta a_1, \qquad o_2 = \zeta a_2$$

From geometry, we have

$$a_1 + m = t + a_2 \rightarrow t = a_1 + m - a_2$$

$$o_1 = t + o_2 \rightarrow t = o_1 - o_2$$

Solving algebraically:

$$a_1 + m - a_2 = o_1 - o_2 = \zeta a_1 - \zeta a_2$$

$$\zeta = \frac{a_1 - a_2 + m}{a_1 - a_2} = 1 + \frac{m}{a_1 - a_2} = 1 + \frac{m}{a_1 - (a_1 + m - t)}$$

$$\zeta = 1 + \frac{m}{t - m}$$

To relate correction factors measured and calculated along the optical axis to the shift applied to the target in FIB view:

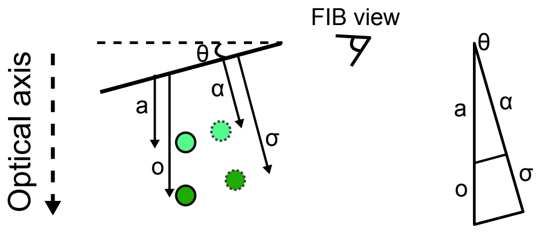


a and o are the actual and apparent depths along the optical axis, $\alpha$ is the apparent target position registered onto the FIB view, and $\sigma$ is the corrected target position in FIB view. Given $o = \zeta a$, from the properties of triangles $\sigma = \zeta\alpha$. This means scaling factors calculated and measured along the optical axis can be directly applied to positions in FIB view, regardless of milling angle.
