## Supplementary material for "High accuracy fluorescence-guided cryo-focused ion beam milling": Optics simulation code: batch_process_vectorial.py

```
"""
This script performs a parallelized parameter sweep to calculate the axial scaling
factor (ζ) for a vectorial PSF model under refractive index mismatch.

It iterates over a range of tilts, depths, NAs, and wavelengths, saving the
results to a CSV file that can be used as a lookup table.
"""

import numpy as np
import pandas as pd
import vectorial_psf_fft as vpsf
import multiprocessing
from itertools import product
from tqdm import tqdm

# The worker function should be in your 'vectorial_psf_fft.py' module
# to ensure it works correctly with multiprocessing.
from vectorial_psf_fft import run_simulation_task

if __name__ == '__main__':
    # --- Define the Parameter Spaces to Sweep ---
    # List all values you want to simulate for each parameter
    tilts_deg = np.arange(0,31,1)
    depths_um = np.arange(0.5,15,0.5)
    nas = np.arange(0.2,1.0,0.1)
    wavelengths_um = np.arange(0.420,0.760,0.050)
    n2=np.arange(1.22,1.48,0.04)

    # --- Create a list of all parameter combinations ---
    param_list = list(product(tilts_deg, depths_um, nas, n2, wavelengths_um))
    
    print(f"Generated {len(param_list)} unique parameter combinations to simulate.")
    print(f"Starting parallel calculation on {multiprocessing.cpu_count()} CPU cores...")

    # --- Run the simulations in parallel ---
    with multiprocessing.Pool() as pool:
        chunksize = max(1, len(param_list) // (multiprocessing.cpu_count() * 16))
        results = list(tqdm(
            pool.imap_unordered(run_simulation_task, param_list, chunksize=chunksize),
            total=len(param_list)
        ))

    # --- Save Results to a Lookup Table ---
    # Create a pandas DataFrame from the list of result tuples
    df = pd.DataFrame(results, columns=['tilt_deg', 'depth_um', 'NA', 'n2', 'wavelength_um', 'zeta'])
    
    # Define the output filename for the data
    output_csv_filename = 'scaling_factor_lookup_table.csv'
    
    # Save the DataFrame to a CSV file
    df.to_csv(output_csv_filename, index=False)

    print("\nCalculation complete.")
    print(f"Lookup table saved to: {output_csv_filename}")
    print("First 5 rows of the table:")
    print(df.head())
```
