## Supplementary material for "High accuracy fluorescence-guided cryo-focused ion beam milling": Optics simulation code: examine_PSF.ipynb

```
# %% [code]
import numpy as np
import pandas as pd
import matplotlib.pyplot as plt
from tifffile import imwrite

import vectorial_psf_fft as vpsf
from vectorial_psf_fft import find_focus_vectorial, run_simulation_task

# %% [code]
def save_psf_tiff_stack(tilt_deg, depth_um, na, wavelength, Nx, Ny):
    # calc geometric parameters
    n_hat = np.array([np.sin(np.deg2rad(tilt_deg)), 0.0, np.cos(np.deg2rad(tilt_deg))])
    dx_um = vpsf.suggest_dx(wavelength, na)

    # calculate pupil plane image
    ctx=vpsf.build_vectorial_pupil_context(n1=1,n2=1.34,NA=na,lam_um=wavelength,zA_um=depth_um,n_hat=n_hat,Nx=Nx,Ny=Ny,dx_um=dx_um)

    # use focus finding routine
    z_star,zeta_axis=vpsf.find_focus_vectorial(zA_um=depth_um,n1=1,n2=1.34,NA=na,lam_um=wavelength,n_hat=n_hat,Nx=Nx,Ny=Ny,z_search_half_width_um=5,Nz=151)
    print(f"Calculated focus at z*={z_star:.3f} um")

    Z=np.arange(z_star-2, z_star+2, 0.2)

    I=[]
    for z in Z:
        # render PSF at each z
        I_z=vpsf.render_plane_vectorial(ctx,z)
        I.append(I_z)
    I=np.array(I)

    # crop I around the center
    crop_radius=40
    I=I[:,Ny//2-crop_radius:Ny//2+crop_radius+1,Nx//2-crop_radius:Nx//2+crop_radius+1]

    imwrite(f'PSF_stack_{tilt_deg}tilt_{depth_um}depth_{na}NA_{int(wavelength*1000)}nm.tif', (I/np.max(I)*65535).astype(np.uint16))

Nx,Ny=513,513

# %% [code]
# easy test case
tilt_deg=0
depth_um=2
na=0.7
wavelength=0.580
save_psf_tiff_stack(tilt_deg=tilt_deg,depth_um=depth_um,na=na,wavelength=wavelength,Nx=Nx,Ny=Ny)

# %% [code]
# a case we care about
tilt_deg=15
depth_um=5
na=0.8
wavelength=0.580
save_psf_tiff_stack(tilt_deg=tilt_deg,depth_um=depth_um,na=na,wavelength=wavelength,Nx=Nx,Ny=Ny)
```
