## Supplementary material for "High accuracy fluorescence-guided cryo-focused ion beam milling": Optics simulation code: vectorial_psf_fft.py

```
# vectorial_psf_fft.py

from __future__ import annotations
import numpy as np
import warnings

__all__ = [
    "suggest_dx",
    "build_vectorial_pupil_context",
    "render_plane_vectorial",
    "find_focus_vectorial",
]

# --------------------------
# Utilities
# --------------------------

def suggest_dx(lam_um: float, NA: float, safety: float = 0.5) -> float:
    return 0.04 # always 40 nm pixels
    # """Suggests lateral sampling Δx (μm) to Nyquist-sample the pupil at a given NA."""
    # return safety * lam_um / (2.0 * max(NA, 1e-9))

def _unit(v: np.ndarray) -> np.ndarray:
    """Normalizes a vector or a batch of vectors to unit length."""
    v = np.asarray(v, dtype=float)
    n = np.linalg.norm(v, axis=-1, keepdims=True)
    n = np.where(n == 0, 1.0, n)
    return v / n

# --------------------------
# Multiprocessing
# --------------------------

def run_simulation_task(params: tuple):
    """
    This is the "worker" function for a single simulation run.
    It unpacks a tuple of parameters, runs the vectorial focus finder,
    and returns the parameters along with the result.
    """
    # Unpack the parameters for this specific simulation task
    tilt_deg, zA_um, NA, n2, lam_um = params

    # --- Define constants for this task ---
    n1= 1.0
    Nx, Ny = 513,513
    n_hat = np.array([np.sin(np.deg2rad(tilt_deg)), 0.0, np.cos(np.deg2rad(tilt_deg))])

    # --- Run the vectorial focus finding simulation ---
    _z_star, zeta_axis = find_focus_vectorial(
        zA_um=float(zA_um),
        n1=n1,
        n2=n2,
        NA=NA,
        lam_um=lam_um,
        n_hat=n_hat,
        Nx=Nx,
        Ny=Ny
    )
    
    # Return the input parameters along with the calculated scaling factor
    return (tilt_deg, zA_um, NA, n2, lam_um, zeta_axis)

# --------------------------
# Core Physics
# --------------------------

def _snell_3d(s1: np.ndarray, n_hat: np.ndarray, n1: float, n2: float) -> np.ndarray:
    """
    Vector Snell's law for a ray originating in medium n2, propagating towards
    an interface with normal n_hat, and refracting into medium n1.

    Args:
        s1: (..., 3) array of unit direction vectors in the immersion medium (n1).
        n_hat: (3,) unit vector normal to the interface, pointing from sample to immersion.
        n1: Refractive index of the immersion medium.
        n2: Refractive index of the sample medium.

    Returns:
        s2: (..., 3) array of corresponding unit direction vectors in the sample medium (n2).
            Rays undergoing TIR are returned as [0, 0, 0].
    """
    n_hat = _unit(n_hat).reshape(3,)
    s1 = np.asarray(s1, dtype=float)
    
    s1_dot_n = np.einsum('...i,i->...', s1, n_hat)
    s1_par = s1 - np.expand_dims(s1_dot_n, -1) * n_hat
    s2_par = (n1 / n2) * s1_par
    
    par2 = np.sum(s2_par**2, axis=-1)
    valid_mask = par2 < 1.0

    s2 = np.zeros_like(s1)
    s2_perp_mag_valid = np.sqrt(1.0 - par2[valid_mask])
    
    # The ray from the emitter travels TOWARDS the interface (+n_hat direction)
    s2[valid_mask] = s2_par[valid_mask] + np.expand_dims(s2_perp_mag_valid, -1) * n_hat

    return s2

# --------------------------
# Vectorial Model Functions
# --------------------------

def build_vectorial_pupil_context(
    *,
    n1: float, n2: float, NA: float, lam_um: float,
    zA_um: float, n_hat: np.ndarray,
    Nx: int = 257, Ny: int = 257, dx_um: float | None = None
) -> dict:
    """
    Builds the vectorial pupil functions for a tilted refractive index
    interface. Uses emission-direction Fresnel coefficients (n2 -> n1)
    which act as angular filters on the light collected by the objective.
    The detection PSF equals the emission PSF by optical reciprocity.
    """
    k0 = 2.0 * np.pi / lam_um
    if dx_um is None:
        dx_um = suggest_dx(lam_um, NA)
    dy_um = dx_um

    # --- 1. Grid & Ray Directions ---
    kx = 2.0 * np.pi * np.fft.fftfreq(Nx, d=dx_um)
    ky = 2.0 * np.pi * np.fft.fftfreq(Ny, d=dy_um)
    KX, KY = np.meshgrid(kx, ky, indexing="xy")
    Kr = np.hypot(KX, KY)

    aperture = (Kr <= k0 * NA)
    
    s1 = np.zeros((Ny, Nx, 3))
    cos_theta1 = np.zeros_like(KX)
    
    kz1 = np.sqrt(np.maximum(0.0, (k0 * n1)**2 - Kr**2))
    s1[aperture, 0] = KX[aperture] / (k0 * n1)
    s1[aperture, 1] = KY[aperture] / (k0 * n1)
    s1[aperture, 2] = kz1[aperture] / (k0 * n1)
    cos_theta1[aperture] = s1[aperture, 2]

    # Refract rays into the sample
    s2 = _snell_3d(s1, n_hat, n1, n2)

    # reject TIR rays
    tir = np.sum(s2**2, axis=-1) < 1e-10
    active = aperture & ~tir

    # --- 2. Phase & Scalar Amplitude ---
    # Phase: Exact spatial phase accumulation exp(i * k2z * zA)
    s2_z = s2[..., 2]
    alpha = k0 * n2 * zA_um * s2_z
    phase_factor = np.exp(1j * alpha)
    
    # Interface normal angles
    cos_a1 = np.abs(np.einsum('...i,i->...', s1, n_hat))
    cos_a2 = np.abs(np.einsum('...i,i->...', s2, n_hat))
    
    safe_cos_theta1 = np.maximum(cos_theta1, 1e-9)

    # Amplitude Correction (Richards-Wolf Framework)
    # Only the Abbe sine condition is required here.
    amp_apodization = 1.0 / np.sqrt(safe_cos_theta1)
    total_scalar_amplitude = amp_apodization
    total_scalar_amplitude[~active] = 0.0

    # --- 3. Local Polarization Basis Vectors ---
    n_hat_3 = n_hat.reshape(1, 1, 3)
    s1_cross_n = np.cross(s1, n_hat_3)
    norm_s1_cross_n = np.linalg.norm(s1_cross_n, axis=-1, keepdims=True)
    norm_s1_cross_n = np.maximum(norm_s1_cross_n, 1e-12)
    
    e_s = s1_cross_n / norm_s1_cross_n
    
    # Handle normal incidence singularity (where s1 || n_hat)
    singularity_mask = (np.linalg.norm(s1_cross_n, axis=-1) < 1e-9)
    e_s[singularity_mask] = np.array([0.0, 1.0, 0.0]) # Arbitrary orthogonal vector
    
    e_p1 = np.cross(e_s, s1) # p-vector in immersion
    e_p2 = np.cross(e_s, s2) # p-vector in sample

    # --- 4. Incident Field (Richards-Wolf Projection) ---
    phi = np.arctan2(KY, KX)
    cos_phi = np.cos(phi)
    sin_phi = np.sin(phi)
    sin_theta1 = np.sqrt(np.maximum(0.0, 1 - cos_theta1**2))
    
    # X-polarized input incident field
    f_xx = cos_phi**2 * cos_theta1 + sin_phi**2
    f_yx = sin_phi * cos_phi * (cos_theta1 - 1)
    f_zx = -cos_phi * sin_theta1

    E_inc_x = np.stack([f_xx, f_yx, f_zx], axis=-1)

    f_xy = f_yx
    f_yy = sin_phi**2 * cos_theta1 + cos_phi**2
    f_zy = -sin_phi * sin_theta1

    E_inc_y = np.stack([f_xy, f_yy, f_zy], axis=-1)

    # --- 5. True Vectorial Transmission ---
    # Decompose into local s and p1 states
    E_s_x = np.einsum('...i,...i->...', E_inc_x, e_s)
    E_p1_x = np.einsum('...i,...i->...', E_inc_x, e_p1)
    
    E_s_y = np.einsum('...i,...i->...', E_inc_y, e_s)
    E_p1_y = np.einsum('...i,...i->...', E_inc_y, e_p1)

    # Fresnel coefficients for EMISSION: n2 -> n1
    # Light from point source in medium 2 transmitting into medium 1
    denom_s = (n2 * cos_a2 + n1 * cos_a1)
    denom_p = (n1 * cos_a2 + n2 * cos_a1)
    ts = np.divide(2 * n2 * cos_a2, denom_s,
                   out=np.zeros_like(denom_s), where=denom_s != 0)
    tp = np.divide(2 * n2 * cos_a2, denom_p,
                   out=np.zeros_like(denom_p), where=denom_p != 0)
    # Reconstruct transmitted field using t_s, t_p, and the new p2 vector
    ts_ext = ts[..., np.newaxis]
    tp_ext = tp[..., np.newaxis]
    
    E_trans_x = (ts_ext * E_s_x[..., np.newaxis]) * e_s + (tp_ext * E_p1_x[..., np.newaxis]) * e_p2
    E_trans_y = (ts_ext * E_s_y[..., np.newaxis]) * e_s + (tp_ext * E_p1_y[..., np.newaxis]) * e_p2

    # --- 6. Final Assembly ---
    # Apply global amplitude and exact phase
    complex_multiplier = (total_scalar_amplitude * phase_factor)[..., np.newaxis]
    
    P_x = complex_multiplier * E_trans_x
    P_y = complex_multiplier * E_trans_y

    beta = -k0 * n1 * cos_theta1

    return dict(
        Px0_x=P_x[..., 0].astype(np.complex128),
        Px0_y=P_x[..., 1].astype(np.complex128),
        Px0_z=P_x[..., 2].astype(np.complex128),
        Py0_x=P_y[..., 0].astype(np.complex128),
        Py0_y=P_y[..., 1].astype(np.complex128),
        Py0_z=P_y[..., 2].astype(np.complex128),
        beta=beta.astype(np.float64),
        Nx=Nx, Ny=Ny, n_hat=n_hat, zA_um=zA_um
    )

def render_plane_vectorial(ctx: dict, z_um: float) -> np.ndarray:
    """
    Renders a 2D intensity image from the vectorial pupil functions at a given z depth.
    """
    propagator = np.exp(1j * ctx["beta"] * float(z_um))
    
    Ex_x = np.fft.fftshift(np.fft.ifft2(ctx["Px0_x"] * propagator))
    Ex_y = np.fft.fftshift(np.fft.ifft2(ctx["Px0_y"] * propagator))
    Ex_z = np.fft.fftshift(np.fft.ifft2(ctx["Px0_z"] * propagator))
    
    Ey_x = np.fft.fftshift(np.fft.ifft2(ctx["Py0_x"] * propagator))
    Ey_y = np.fft.fftshift(np.fft.ifft2(ctx["Py0_y"] * propagator))
    Ey_z = np.fft.fftshift(np.fft.ifft2(ctx["Py0_z"] * propagator))
    
    I_x = np.abs(Ex_x)**2 + np.abs(Ex_y)**2 + np.abs(Ex_z)**2
    I_y = np.abs(Ey_x)**2 + np.abs(Ey_y)**2 + np.abs(Ey_z)**2
    
    return I_x + I_y

def find_focus_vectorial(
    zA_um: float,
    n1: float, n2: float, NA: float, lam_um: float, n_hat: np.ndarray,
    Nx: int = 257, Ny: int = 257,
    z_search_half_width_um: float = 5.0,
    Nz: int = 151,
    plot_fit: bool = False
) -> tuple[float, float]:
    """
    Robustly finds the focus for the vectorial model by fitting a parabola 
    to the axial intensity profile in a fixed 11-point window around the discrete maximum.
    """
    dx_um = suggest_dx(lam_um, NA)
    ctx = build_vectorial_pupil_context(
        n1=n1, n2=n2, NA=NA, lam_um=lam_um, zA_um=zA_um, n_hat=n_hat,
        Nx=Nx, Ny=Ny, dx_um=dx_um
    )

    z_center_paraxial = zA_um * (n1 / n2)
    z_grid_um = np.linspace(
        z_center_paraxial - z_search_half_width_um,
        z_center_paraxial + z_search_half_width_um,
        Nz
    )
    
    axial_profile = np.empty(Nz, dtype=float)
    
    for k, z in enumerate(z_grid_um):
        I = render_plane_vectorial(ctx, z)
        axial_profile[k] = I.max()

    # --- Parabola Fitting Logic ---
    
    # 1. Find the raw discrete maximum
    idx_max = int(np.argmax(axial_profile))
    z_max_discrete = z_grid_um[idx_max]
    
    # 2. Grab 5 points on either side (up to 11 points total)
    idx_start = max(0, idx_max - 5)
    idx_end = min(Nz, idx_max + 6)
    
    z_window = z_grid_um[idx_start:idx_end]
    I_window = axial_profile[idx_start:idx_end]
    
    # 3. Fit 2nd order polynomial: I(z) = A*z^2 + B*z + C
    try:
        A, B, C = np.polyfit(z_window, I_window, 2)
        
        # The peak of A*z^2 + B*z + C is at z = -B / (2A)
        # Check that it's a downward-opening parabola
        if A < 0:
            z_star_fit = -B / (2.0 * A)
            
            # Sanity check: Ensure the fitted peak isn't wildly outside our window
            # (e.g., if the data is so asymmetric that the parabola apex flies off)
            if z_window[0] <= z_star_fit <= z_window[-1]:
                z_star = z_star_fit
            else:
                z_star = float(z_max_discrete)
        else:
            # Opens upward or is flat; fallback to discrete
            z_star = float(z_max_discrete)
            
    except Exception as e:
        warnings.warn(f"Parabola fit failed ({e}). Falling back to discrete max.")
        z_star = float(z_max_discrete)
        A = 1 # Force skip plot fitting

    # --- Diagnostic Plotting ---
    if plot_fit:
        import matplotlib.pyplot as plt
        plt.figure(figsize=(8, 5))
        
        plt.plot(z_grid_um, axial_profile, 'k.-', label='Raw Axial Profile', alpha=0.6)
        plt.plot(z_window, I_window, 'ro', label='Fitting Window (11 pts)', markersize=6)
        
        if A < 0:
            # Plot the smooth parabolic fit over the window range
            z_dense = np.linspace(z_window[0], z_window[-1], 100)
            plt.plot(z_dense, A * z_dense**2 + B * z_dense + C, 'b-', linewidth=2, label='Fitted Parabola')
            plt.axvline(z_star, color='b', linestyle=':', label=f'Found Focus: z* = {z_star:.4f}')
            plt.title("Diagnostic: Parabola Fit Successful")
        else:
            plt.axvline(z_max_discrete, color='r', linestyle=':', label=f'Fallback: z* = {z_max_discrete:.4f}')
            plt.title("Diagnostic: Parabola Fit FAILED / REJECTED")
            
        plt.xlabel('Nominal Focus z (μm)')
        plt.ylabel('Peak Intensity')
        plt.legend()
        plt.grid(True, alpha=0.3)
        plt.tight_layout()
        plt.show()

    # Depth scaling
    zeta_axis = zA_um / z_star if z_star != 0 else np.inf
    
    return z_star, zeta_axis

## centroid version
# def find_focus_vectorial(
#     zA_um: float,
#     n1: float, n2: float, NA: float, lam_um: float, n_hat: np.ndarray,
#     Nx: int = 257, Ny: int = 257,
#     z_search_half_width_um: float = 5.0,
#     Nz: int = 151
# ) -> tuple[float, float]:
#     """
#     Robustly finds the focus for the vectorial model using the axial centroid.
#     """
#     dx_um = suggest_dx(lam_um, NA)
#     ctx = build_vectorial_pupil_context(
#         n1=n1, n2=n2, NA=NA, lam_um=lam_um, zA_um=zA_um, n_hat=n_hat,
#         Nx=Nx, Ny=Ny, dx_um=dx_um
#     )

#     z_center_paraxial = zA_um * (n1 / n2)
#     z_grid_um = np.linspace(
#         z_center_paraxial - z_search_half_width_um,
#         z_center_paraxial + z_search_half_width_um,
#         Nz
#     )
    
#     axial_profile = np.empty(Nz, dtype=float)
    
#     for k, z in enumerate(z_grid_um):
#         I = render_plane_vectorial(ctx, z)
#         axial_profile[k] = I.max()

#     axial_profile_baselined = axial_profile - np.min(axial_profile)
#     total_intensity_sum = np.sum(axial_profile_baselined)
#     if total_intensity_sum > 1e-9:
#         z_star = np.sum(z_grid_um * axial_profile_baselined) / total_intensity_sum
#     else:
#         z_star = z_grid_um[Nz // 2]

#     # Since zA_um is now defined along the optical axis, no cos(tau) projection is needed.
#     zeta_axis = zA_um / z_star if z_star != 0 else np.inf
    
#     return z_star, zeta_axis
```
